# Model of ocular surface ion and water transport predicts efficacy of dry eye therapeutics targeting epithelial transport and tear fluid dynamics

**DOI:** 10.1101/2025.11.17.688783

**Authors:** Varun Verma, Ethan S. Lindgren, Marc H. Levin, Onur Cil, Lukmanee Tradtrantip, Rongshan Yan, Neel D. Pasricha, Alan S. Verkman

## Abstract

The composition and volume of tear fluid lining the ocular surface are governed by the rates of lacrimal gland secretion, tear film evaporation, nasolacrimal drainage, and epithelial ion and water transport. Tear fluid hyperosmolality and reduced volume are key drivers of dry eye disease (DED) pathogenesis. We constructed a mathematical model to compute the composition and volume of tear fluid and epithelial cell compartments, with transport parameters specified for the mouse eye from published data and in vivo measurements of ocular surface potential differences. The model accounted for transcellular and paracellular transport across the epithelia under open-circuit conditions utilizing flux equations for individual transport mechanisms, with mass balance constraints on solute and water content in cytoplasm and tear fluid. Under DED conditions established by reduced lacrimal secretion and increased evaporation, the model predicted the efficacy of currently available DED therapies including punctal plugs, humidification goggles, lacrimal gland stimulation, and artificial tears eye drops. The model also predicted the limited efficacy of anti-absorptive and pro-secretory drugs targeting epithelial ion transporters, and the high efficacy of targeting epithelial water permeability or paracellular ion permeability. The modeling herein provided quantitative predictions to prioritize novel targets for DED and drive the development of new therapies.

**Author Summary:** Dry eye disease (DED) affects billions of adults worldwide, but a full picture is lacking of how the tear film becomes abnormally thin and hyperosmolar. The computer model built here links four processes – tear production by the lacrimal gland, tear fluid evaporation, tear drainage through tear ducts, and transport of solutes and water across eye surface epithelial cells – to predict the thickness and composition of the tear film in various conditions. Model parameters were selected using published data and electrical measurements of voltage changes across the ocular surface produced by ion transport. The model predicted that existing therapies, such as punctal plugs, moisture goggles, or stimulating tear production, can substantially increase tear thickness or lower saltiness. The model also predicted limited efficacy of drug therapies in current development that target ion transport, and identified epithelial cell water transport and paracellular ion permeability as novel targets for DED treatment. By making ocular surface transport mechanisms explicit and testable, our work offers a roadmap for development of new therapies that restore a healthy tear film, a major unmet medical need.

## Introduction

The volume, osmolality, and composition of the tear film are key determinants of corneal and conjunctival health [1]. Abnormalities in tear fluid are associated with ocular pathology, including dry eye disease (DED), in which tear fluid hyperosmolality and reduced volume drive inflammation and consequent pain and tissue injury [2]. DED is widespread, affecting 1 out of ∼10 adults over the age of 50 in the United States, with an estimated global prevalence of 30% [1,3]. Tear fluid composition and volume are regulated by coordinated lacrimal gland secretion, tear fluid evaporation, nasolacrimal drainage, and transepithelial solute and water transport across the cornea and conjunctiva [2,4]. Impaired lacrimal gland secretion and excessive evaporative water loss underlie aqueous-deficient and evaporative dry eye, respectively [2]. Current therapies targeting the determinants of tear fluid composition and volume include chemical or electrical stimulation to enhance lacrimal gland secretion [5,6], lipid-containing eye drop formulations and moisture goggles to reduce evaporation [7,8], and punctal occlusion to reduce nasolacrimal drainage [9]. Despite these options, DED remains inadequately controlled in approximately 50% of patients [10].

The ocular surface epithelium, consisting of cornea and conjunctiva, makes important yet comparatively understudied contributions to tear fluid homeostasis. The ocular surface epithelium can accomplish net fluid absorption or secretion depending on electrochemical driving forces and transporter activities [4]. Classic studies implicate the epithelial Na^+^ channel (ENaC) in fluid absorption, and cAMP- and Ca^2+^-activated Cl^−^ channels, including the cystic fibrosis transmembrane conductance regulator (CFTR), in fluid secretion [11–13]. Ocular surface epithelial cells also express K^+^ channels, the Na^+^/K^+^/2Cl^−^ cotransporter NKCC1, Na^+^-coupled and Na^+^-independent glucose and amino acid transporters, aquaporins (AQPs), a Na^+^/K^+^-ATPase, and additional transport pathways; paracellular ion and solute movement also contributes to net transepithelial transport [4]. Small-molecule therapeutics in development for DED that target epithelial ion transport include inhibitors of transcellular Na^+^ absorption [14], and activators of transcellular Cl^−^ secretion acting on CFTR [15] or Ca^2+^-activated Cl^−^ channels [16].

Herein, we develop and apply a quantitative model to predict the composition and volume of the tear fluid and epithelial cell compartments. The model provides a rigorous framework to quantify the dependence of tear fluid properties on lacrimal secretion, tear film evaporation, nasolacrimal drainage, and epithelial transport. An important goal of the modeling was to predict the efficacy of approved and investigational dry eye therapies, and to identify novel targets for drug development. Model parameters were selected for the mouse eye, leveraging existing literature [17–20] and in vivo ocular surface potential difference (OSPD) measurements done as part of this study. Although the model formulation and numerical solution are complex, the model yielded clear-cut insights into DED mechanisms and drug effects, including the identification of previously unexplored targets for dry eye drug development.

## Results

### Model formulation

As diagrammed in Fig. 1, the model comprises an epithelial cell monolayer in contact with a tear fluid layer, both assumed to be well-mixed. The cytoplasm and tear fluid compartments contain the primary solutes Na^+^, K^+^, Cl^-^ and glucose, with concentrations denoted [X]_cell_ and [X]_tear_, respectively, and aqueous volumes described by tear heights h_cell_ and h_tear_. Tear fluid volume and composition are determined by the rates of lacrimal secretion, tear fluid evaporation, nasolacrimal drainage and epithelial transport. Fluid enters the tear film by lacrimal secretion defined by solute and water fluxes, J_s_^lac^ and J_v_^lac^, respectively, and fluid exits by nasolacrimal drainage, J_s_^drain^ and J_v_^drain^ ; evaporation removes solute-free water from the tear fluid, J_v_^evap^.

**Figure 1.**
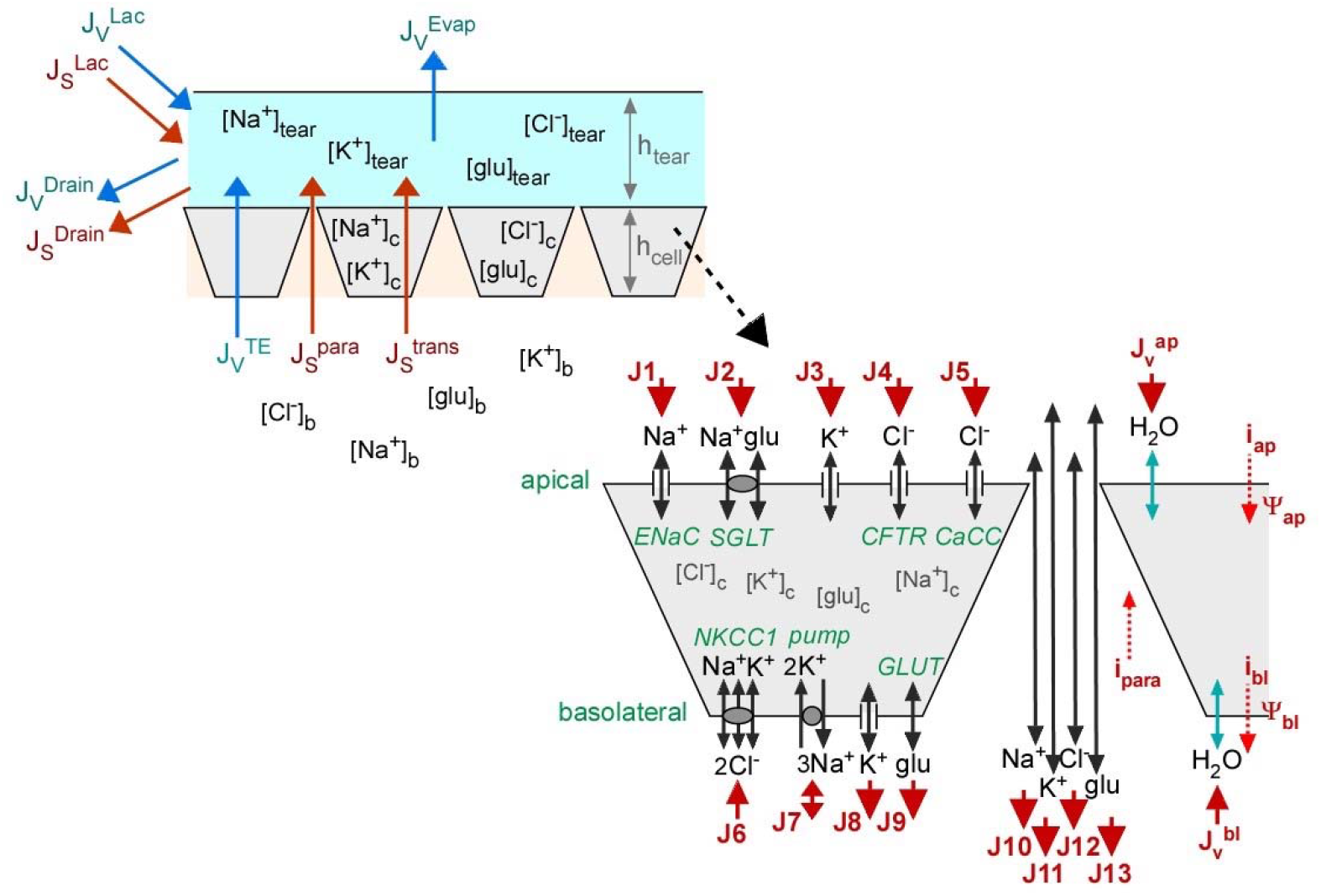
Schematic model of tear fluid volume and osmolality. (left) Epithelial cell layer in contact with the tear film, showing solute (J_s_) and volume (J_v_) fluxes for lacrimal gland fluid secretion, tear film evaporation, nasolacrimal drainage, and epithelial transport. See Table 1 for definitions of model symbols. (right) Epithelial transporters included in the model. Apical and basolateral membrane transporters shown, as well as paracellular transport. Solute fluxes (J1-J13) indicated with red arrows denoting directions of positive flux. For J7, the pump operates exclusively in a Na^+^ extrusion mode. Apical and basolateral membrane water fluxes (J_v_) shown on the right, along with total ionic currents (i_ap_, i_bl_, i_para_) and membrane potentials (ψ_ap_, ψ_bl_).

Transepithelial transport occurs via transcellular and paracellular pathways, the former involving transport across apical (tear film-facing) and basolateral (stroma/underlying tissue-facing) epithelial cell plasma membranes. The epithelial model (Fig. 1, right) includes apical ENaC, CFTR/CaCC, and K^+^ channels; basolateral K^+^ channels; an apical SGLT; a basolateral NKCC1; an electroneutral glucose transporter; and a basolateral 3Na^+^/2K^+^-ATPase (pump) that energizes transport. Osmotically driven water flow crosses membranes via AQP and AQP-independent pathways. Paracellular ion and glucose permeation occurs as well. The figure also shows total ionic currents (red dashed arrows; i_ap_, i_bl_, i_para_) and cell plasma membrane potentials (ψ_ap_, ψ_bl_).

**Table 1.** Model symbols and definitions.

|  |  |
| --- | --- |
| $[X]_{\text{tear}}, [X]_{\text{cell}}, [X]_{\text{bl}}, [X]_{\text{lac}}$ | Concentration of solute X (tear, cell, basolateral, lacrimal) (mM) |
| $\phi_X$ | Osmotic coefficient of solute X in cytoplasm |
| $\beta_X$ | Osmotic coefficient of solute X in extracellular space |
| $P_f^{\text{ap}}, P_f^{\text{bl}}$ | Apical and basolateral osmotic water permeability coefficient (cm/s) |
| $V_w$ | Partial molar volume of water ($\text{cm}^3/\text{mol}$) |
| J1-J9 | Membrane transporter solute fluxes / turnover rates ($\mu\text{mol}/\text{cm}^2\text{s}$) |
| J10-J13 | Paracellular solute fluxes ($\mu\text{mol}/\text{cm}^2\text{s}$) |
| P1-P13 | Transporter permeability coefficients (as defined in Eqns. 1-13) |
| $\psi_{\text{ap}}, \psi_{\text{bl}}, \psi_{\text{te}}$ | Membrane (apical, basolateral) and transepithelial potentials (mV) |
| F/RT | F, Faraday constant; R, gas constant; T, absolute temperature ($\text{mV}^{-1}$) |
| $[X]_{\text{lac}}$ | Concentration of solute X in fluid secreted by lacrimal gland (mM) |
| $\text{Osm}_{\text{tear}}, \text{Osm}_{\text{cell}}, \text{Osm}_{\text{blood}}, \text{Osm}_{\text{lac}}$ | Osmolalities (tear fluid, cytoplasm, blood, lacrimal fluid) (mOsm) |
| $h_{\text{tear}}, h_{\text{cell}}$ | Compartment heights (tear fluid, cell) ($\mu\text{m}$) |
| $J_v^{\text{ap}}, J_v^{\text{bl}}, J_v^{\text{te}}$ | Volume fluxes (apical, basolateral, transepithelial) ($\text{cm}^3/\text{cm}^2\text{s}$) |
| $J_v^{\text{lac}}, J_v^{\text{evap}}, J_v^{\text{drain}}$ | Volume fluxes (lacrimal, evaporative, drain) ($\text{cm}^3/\text{cm}^2\text{s}$) |
| $J_{\text{osm}}^{\text{ap}}, J_{\text{osm}}^{\text{bl}}, J_{\text{osm}}^{\text{te}}$ | Osmolar fluxes (apical, basolateral, transepithelial) ($\mu\text{osmol}/\text{cm}^2\text{s}$) |
| $J_{\text{osm}}^{\text{lac}}, J_{\text{osm}}^{\text{cell}}$ | Osmolar fluxes (lacrimal, cytoplasm) ($\mu\text{osmol}/\text{cm}^2\text{s}$) |
| $J_X^{\text{tear}}, J_X^{\text{cell}}$ | Solute fluxes (out of tear fluid, into cell) ($\mu\text{mol}/\text{cm}^2\text{s}$) |
| $J_X^{\text{ap}}, J_X^{\text{bl}}, J_X^{\text{lac}}, J_X^{\text{drain}}$ | Solute fluxes (apical, basolateral, lacrimal, drain) ($\mu\text{mol}/\text{m}^2\text{s}$) |
| $\Delta t$ | Time step (s) |
| $i_{\text{ap}}, i_{\text{bl}}, i_{\text{para}}$ | Currents (apical, basolateral, paracellular) ($\mu\text{A}/\text{cm}^2$) |
| $K_i^X$ | Saturation constant for solute X, transporter i (mM) |
| a, b | Coefficients for Na/K pump ($b=1-a\psi_{\text{bl}}F/RT$) |

The model computes tear film and cell cytoplasm variables (solute concentrations, osmolalities, membrane potentials, layer heights) for a specified set of input parameters that can be modified at defined times. Model parameters are defined in Table 1 and parameter values used in the computations are listed in Supplementary Table S1. Fig. 2 shows the computational approach to determine the kinetics of tear film and cellular variables in response to changes in input parameters. Briefly, at each time point cell membrane potentials are computed iteratively from ion fluxes, subject to open-circuit electroneutrality requirements (equal apical, basolateral and paracellular ion currents). Tear fluid and cell heights are determined from osmolar and consequent volume fluxes arising from epithelial transport, as well as lacrimal gland secretion, tear film evaporation, and nasolacrimal drainage. Tear fluid and cytoplasmic solute concentrations are then computed. The process is repeated for the specified total simulation time with 0.1 s time steps. Predicted effects of experimental maneuvers, such as addition of epithelial transport modulators or reducing evaporation rate, are modeled by changing input parameters at specified times.

**Figure 2.**
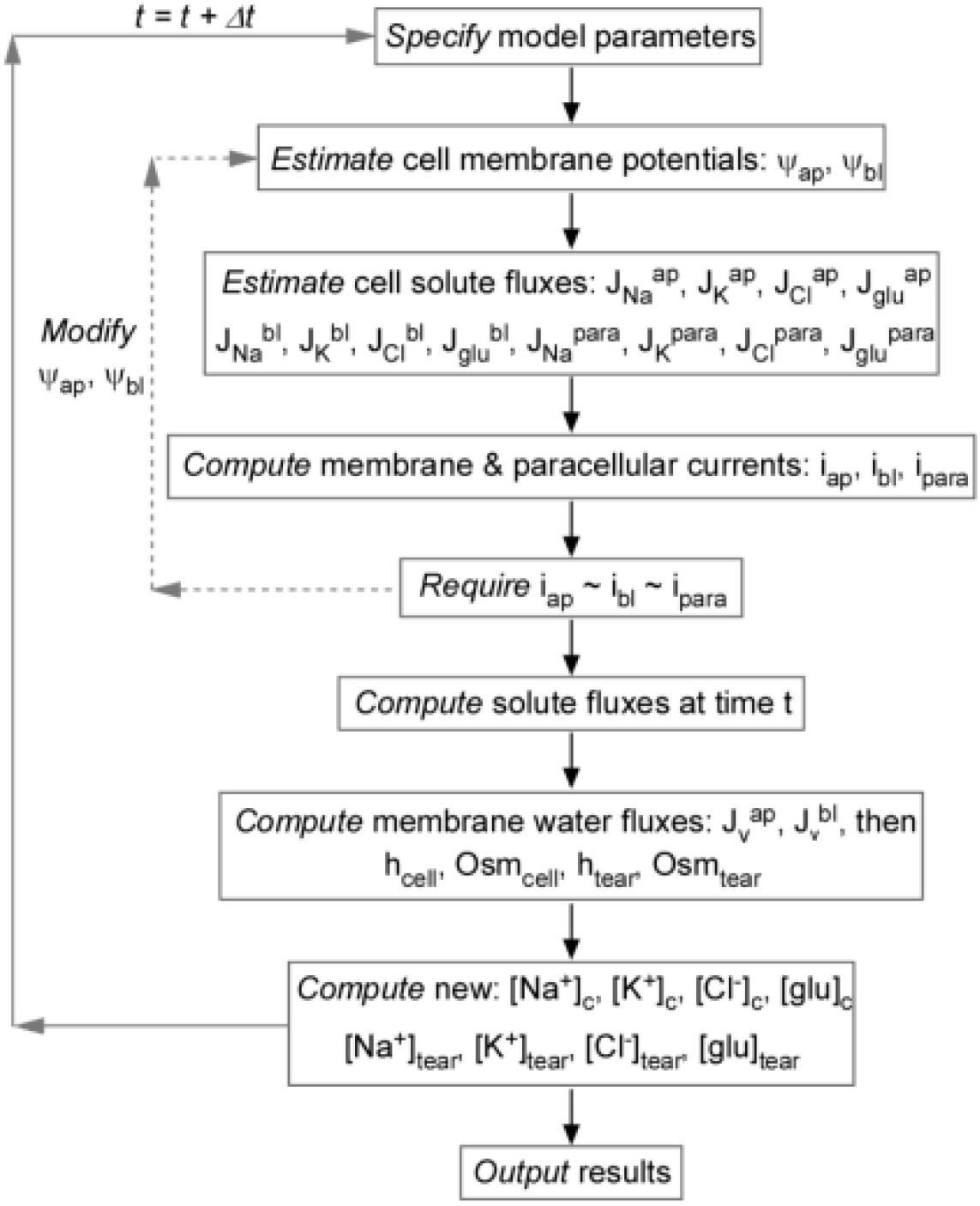
Numerical solution of the tear fluid model. The time course of model variables is computed in response to changes in epithelial solute concentrations or permeability parameters or rates of lacrimal gland secretion, evaporation or nasolacrimal drainage. See text for explanations.

Thirteen epithelial solute fluxes J1–J13 (red arrows in Fig. 1 denote positive direction) comprise 5 apical, 4 basolateral, and 4 paracellular pathways, as adapted from ref. [17] with minor modifications. Goldman–Hodgkin–Katz relations are used to describe channel/leak fluxes, solute carrier equations include saturability, and the basolateral Na^+^/K^+^ ATPase provides active transport.

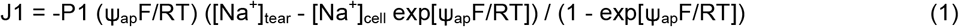

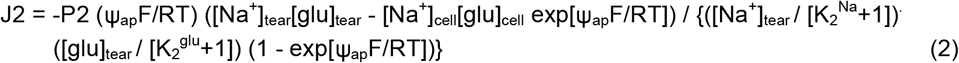

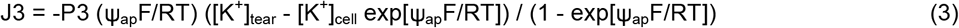

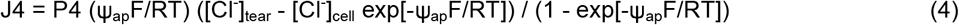

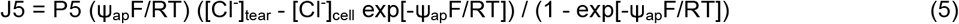

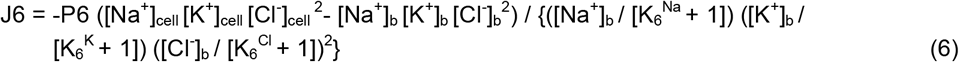

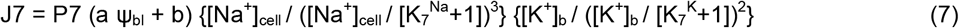

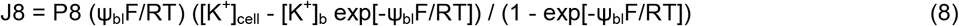

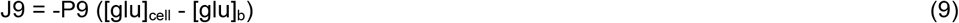

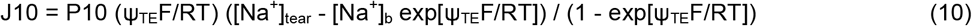

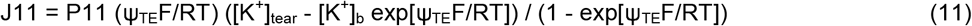

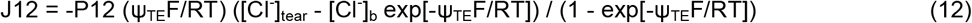

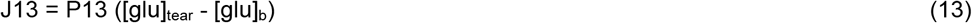

Open-circuit electroneutrality requires,

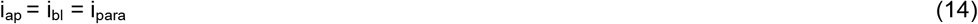

where,

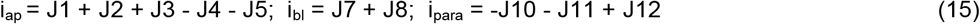

Starting with initial guesses for membrane potentials and solute concentrations (tear fluid and cytoplasm), ψ_ap_ and ψ_bl_ are adjusted to give electroneutrality using 2-variable Newton iteration (Jacobian of i_ap_ - i_bl_ and i_ap_ - i_para_ with respect to ψ_ap_ and ψ_bl_).

Following the electroneutrality computation at time t, cell and tear fluid heights (volume per area), are computed from osmotic water fluxes and, for the tear fluid, the rates of volume influx from lacrimal secretion and volume loss from evaporation and nasolacrimal drainage,

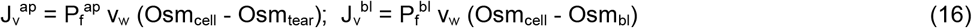

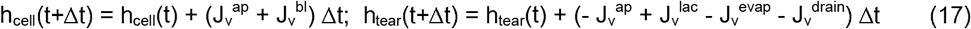

Osmolalities of the tear fluid and cytoplasm at time t + Δt are computed from solute fluxes,

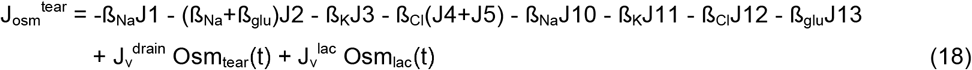

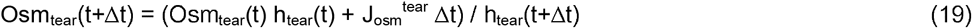

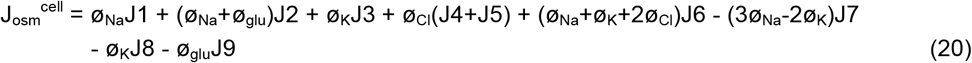

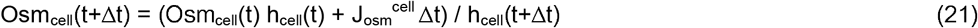

Lastly, solute concentrations at time t + Δt are computed from individual solute fluxes. In the tear fluid compartment,

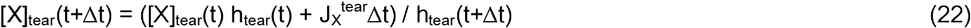

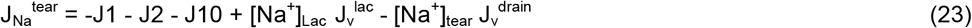

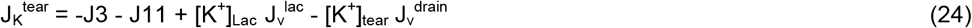

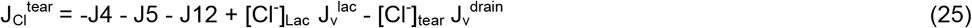

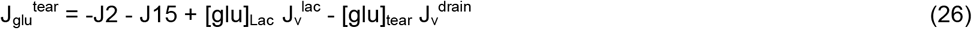

In the cytoplasmic compartment,

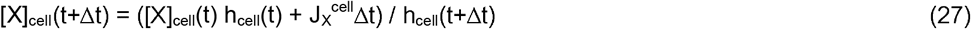

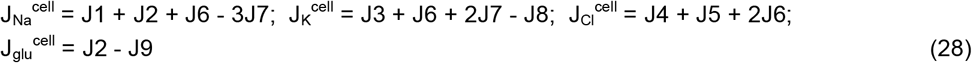

Equations 1–28 are solved at successive time steps to obtain the time dependence of the computed variables. The simulations were implemented in Python. Results were insensitive to time step size (0.005-0.1 s), floating-point precision, and initial guesses for membrane potentials and concentrations, indicating robust numerical behavior. To validate the kinetic simulations, we also implemented a steady-state solver formulated as a 12-variable nonlinear least-squares problem for Eqs. S1–S12 (see Supplementary Data), enforcing electroneutrality and zero net flux of each solute and water in the cellular and tear fluid compartments. Kinetic simulations carried out for 7,000 seconds agreed with steady-state computations to within 1%, confirming computational accuracy.

### Parameter selection

Epithelial transport parameters were adapted from Levin et al [17], with membrane and paracellular permeabilities refined using OSPD data in which open-circuit potential differences were measured across the mouse ocular surface epithelium in vivo during continuous perfusion with solutions contacting the ocular surface. Representative OSPD curves are shown in Fig. 3. The baseline fluid composition for initial perfusions was chosen to approximate reported tear fluid composition and osmolality. The maneuvers used to probe epithelial transport included ion (Na^+^, K^+^, Cl^-^) substitutions and application of transport inhibitors (amiloride for ENaC, CFTR_inh_-172 for CFTR, BaCl_2_ for K^+^ channels) and activators (cAMP agonist forskolin to activate CFTR and basolateral K^+^ channels, ATP to activate CaCC and K^+^ channels). As adapted from prior OSPD measurements in different species including humans [15,17,21], the mouse ocular surface was exposed to serial additions of amiloride, Cl^-^ free solution, cAMP activator, CFTR inhibitor, and ATP (Fig. 3A). Then, in order to assess composite transcellular/paracellular versus paracellular ion transport pathways, OSPD changes were measured upon removal and reintroduction of individual ions in the absence and the presence of transport inhibitors (OSPD curves on left vs. right in Fig. 3B-D). Finally, in order to quantify total paracellular ionic conductance a study was done in which Na^+^, K^+^ and Cl^-^ were replaced with sucrose, without and with transport inhibitors (Fig. 3E). Because of intrinsic biological variability and technical challenges in these in vivo OSPD measurements, baseline OSPD values and changes in OSPD with various maneuvers varied between mice. For parameter selected, averaged experimental values for each maneuver were used, as summarized in Supplementary Table S2.

**Figure 3.**
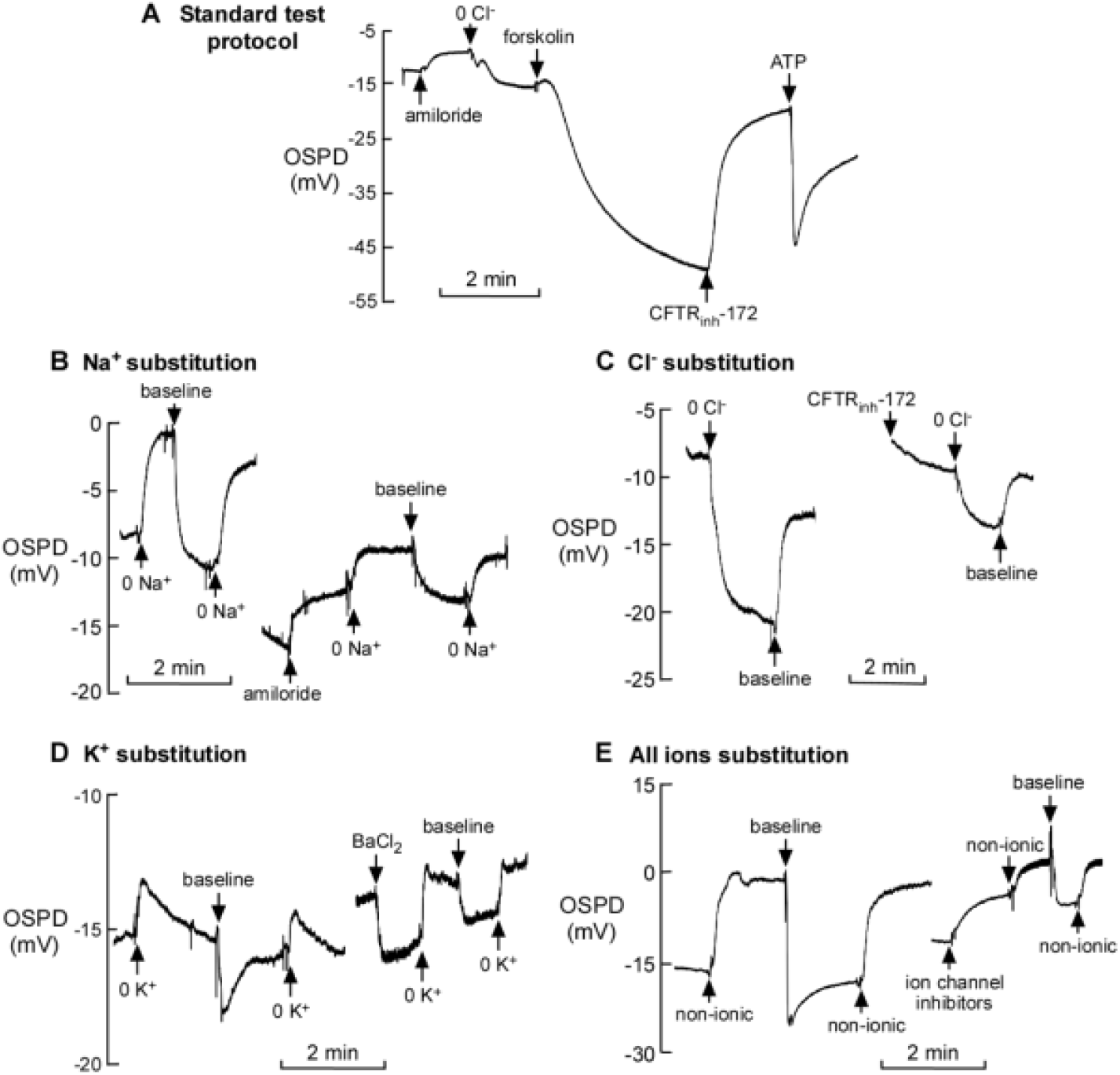
OSPD measurements in live mice. The ocular surface was perfused with solutions of specified ionic compositions (see Methods) with and without selective transport modulators, as indicated. Standardized protocol (A) showing serial responses to amiloride (ENaC inhibitor), Cl^-^ substitution, forskolin (CFTR / K^+^ channel activator), CFTR inhibitor, and ATP (CaCC / K^+^ channel activator). Ionic substitution experiments shown for Na^+^ (B), Cl^-^ (C), K^+^ (D), and all three ions together (E), with ion substitutions done in the absence (left) and presence (right) of transport inhibitors. Representative experiments shown; OSPD changes are summarized in Supplemental Table S2. Concentrations: amiloride, 100 µM; CFTR_inh_-172, 10 µM; forskolin, 10 µM; Ba^2+^, 5 mM; ATP, 100 µM.

Paracellular ion permeabilities were deduced by minimizing the differences between measured and computed OSPD across the panel of experimental maneuvers using our previously reported one-compartment model [17]. Using the deduced parameter set (Supplementary Table S1), there was generally good agreement between experimental and computed OSPD values (Supplementary Table S2) as well as baseline OSPD, ion concentrations, and membrane potentials (Supplementary Table S3).

The set of refined epithelial parameters was used in modeling the tear film, which also required specification of parameters for lacrimal gland secretion, tear film evaporation, and nasolacrimal drainage. The relations and parameters defining these processes are discussed in the Supplementary Data and summarized in Supplementary Figs. S1-S3. Briefly, evaporative water loss from the tear fluid increases as tear fluid volume is reduced because break-up and thinning of the superficial lipid layer lowers the evaporative barrier and exposes more aqueous surface [2,22]. For lacrimal gland secretion, reported measurements indicate mildly hyperosmolar fluid at low secretion rates and near-isosmolar fluid at higher secretion rates [23]. For nasolacrimal drainage, loss of bulk tear fluid increases with tear volume as limited by the flow capacity of nasolacrimal ducts [24].

### Model behavior

Fig. 4 shows the effects of changing evaporation, nasolacrimal drainage and lacrimal secretion on the principal tear variables, tear fluid height and osmolality, with all variables listed in Supplementary Tables S5-S7 for selected evaporation, drainage and lacrimal secretion rates. Doubling evaporation produced a substantial, 16 mOsm, increase in tear osmolality with minimal reduction in tear height. Evaporation causes solute-free water loss, which, by increasing tear fluid osmolality, drives transepithelial water secretion.

**Figure 4.**
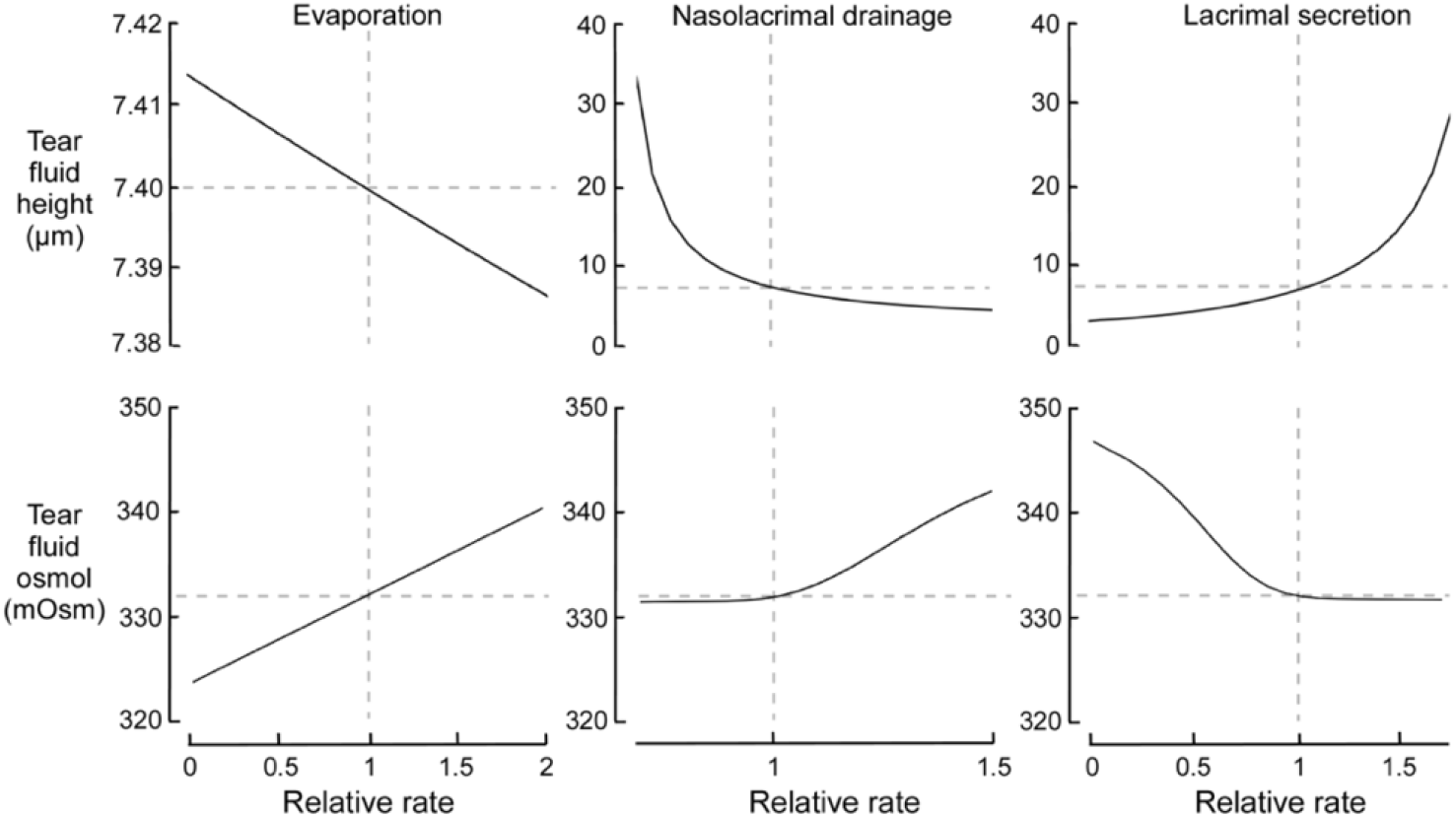
Dependence of tear fluid height and osmolality on evaporation, nasolacrimal drainage and lacrimal secretion. Steady-state computations done under baseline conditions (parameters listed in Supplementary Table S1 and Figs. S1-S3), keeping parameters unchanged except for tear film evaporation (left), nasolacrimal drainage (middle), or lacrimal secretion (right). See Tables S5-S7 in Supplementary Data for associated membranes potentials, concentrations, and fluxes. Dashed horizontal grey lines indicate baseline h_tear_ (7.4 µm) and Osm_tear_ (332 mOsm); dashed vertical grey lines indicate baseline rate (unity relative rate).

Reduced nasolacrimal drainage (relative rate < 1) strongly increased tear film height, predicting epiphora (tear overflow) with full punctal occlusion in healthy, non-DED mice, consistent with human studies [9], but with minimal changes in tear fluid osmolality. Increased evaporation (relative rate > 1) produced a significant, 10 mOsm, increase in tear osmolality with a relatively small effect on tear film height.

Similar predictions were seen with increased lacrimal gland secretion (relative rate > 1), which produced increased tear film height with relatively small changes in osmolality. Reduced lacrimal gland secretion (relative rate < 1) produced relatively small reductions in tear fluid height and more substantial increases in osmolality. These findings align with the model descriptions of tear film evaporation, drainage and lacrimal secretion shown in Supplementary Figs. S1-S3. A general observation from these and subsequent computations is the rough orthogonality of lacrimal secretion and nasolacrimal drainage in determining tear fluid height vs. that of evaporation and transepithelial osmosis in determining tear fluid osmolality. However, the precise relationship among these phenomena is complex as formally described by model equations 1-28 and the tear film evaporation, drainage, and lacrimal secretion relations in Supplementary Figs. S1-S3.

Fig. 5 shows the effects of varying selected epithelial transport parameters on steady-state tear fluid osmolality and height. Changing apical K^+^ conductance (P3), Cl^−^ conductance (P5), basolateral K^+^ conductance (P8), or glucose transport (P9) had minimal effects on tear fluid parameters (not shown). More substantial increases in tear film height were predicted with modulation of ENaC (P1), CFTR (P4), NKCC1 (P6), paracellular Na^+^/K^+^/Cl^−^ conductances (P10-P12), and apical/basolateral water permeability (P_f_^ap^/P_f_^bl^). Notably, ENaC inhibition produced a substantial increase in tear height, while relatively modest changes in tear film osmolality and height were seen with increased CFTR or NKCC1 permeabilities. Activation of the 3Na^+^/2K^+^ ATPase (P7) or inhibition of the Na^+^/glucose symporter (P2) produced lesser but significant changes following the same patterns as seen for ENaC, NKCC1 or CFTR modulation (not shown).

**Figure 5.**
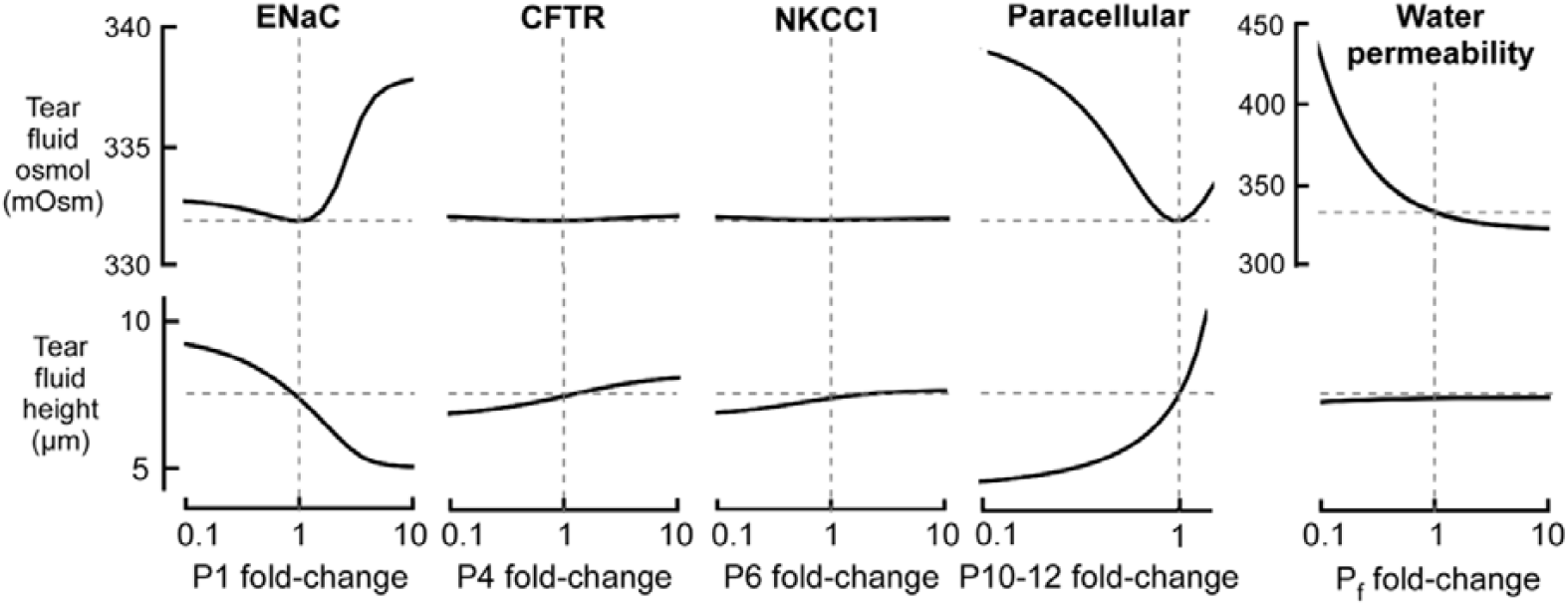
Dependence of tear film height and osmolality on epithelial parameters. Steady-state computations done under baseline conditions (parameters listed in Supplementary Table S1 and Figs. S1-S3), keeping parameters unchanged except for permeabilities associated with ENaC (P1), CFTR (P4), NKCC1 (P6), paracellular permeability (P10-13, changes made in parallel), and water permeability (P_f_^ap^ and P_f_^b^, changes made in parallel). Dashed horizontal grey lines indicate baseline h_tear_ (7.4 µm) and Osm_tear_ (332 mOsm); dashed vertical grey lines indicate baseline rate (unity relative rate).

A substantial increase in tear film height was predicted with a relatively small, 1.5-fold increase in paracellular ion conductances, which promotes paracellular ion and consequent transepithelial water secretion into the tear film. Remarkably, increasing cell water permeability strongly reduced tear film osmolality with minimal effects on height, which is a consequence of increased transepithelial osmotic water secretion into the tear film.

### Tear film dynamics

Modeling tear film dynamics in response to clinically relevant maneuvers is shown in Fig. 6, as applicable, for example, to the onset of action of dry eye therapeutics or environmental changes in ambient humidity. Altered evaporation produced transient changes in tear fluid height with return to baseline in tens of minutes, but with rapid and sustained changes in tear fluid osmolality. The directions of these effects were as expected. Increased evaporation, as would occur in transiting to a dry air environment, increased tear fluid osmolality and transiently reduced tear film height. Reduced tear film evaporation, as would occur with protective goggles or lipid-containing eye drop formulations, produced a significant, sustained reduction in tear fluid osmolality and transient increase in height. Reduced nasolacrimal drainage or increased lacrimal secretion produced relatively slow increases in tear film height, but with rapid, small reductions in tear film osmolality. The relatively slow response in tear film height is a consequence of relatively slow kinetics of lacrimal fluid entry into and nasolacrimal fluid drainage out of the tear film, compared to the more rapid changes in tear fluid osmolality afforded by evaporative water loss and transepithelial osmosis.

**Figure 6.**
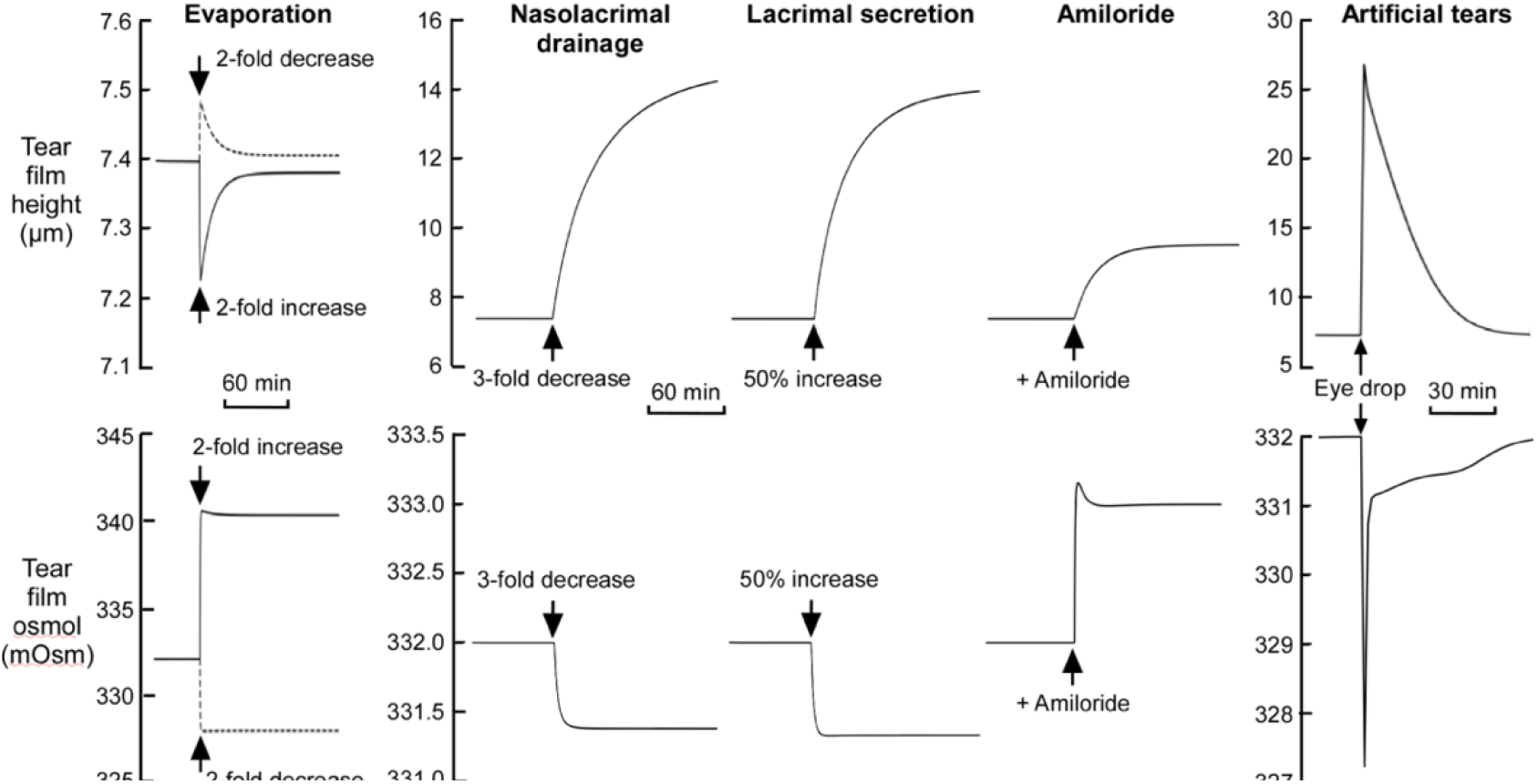
Kinetics of tear fluid height and osmolality in response to clinically relevant changes in model parameters. Computations done for indicated changes (arrows) in evaporation rate, nasolacrimal drainage, lacrimal secretion, addition of ENaC inhibitor amiloride (to give 100% inhibition), and addition of a 2 µl artificial tears eye drop (325 mOsm, Na□ 112 mM, K□ 3.2 mM, Cl□ 84 mM).

Modeling the pharmacodynamics of amiloride, inhibition of apical ENaC Na□ conductance produced a smaller, albeit significant, increase in tear fluid height and interestingly a small rapid increase in tear fluid osmolality. This occurs because ENaC inhibition reduces Na□ absorption and consequent net ion retention in the tear film without direct effects on water evaporation or transepithelial transport. The steady-state following amiloride application was achieved in ∼60 minutes. These results are relevant to the clinical testing of ENaC inhibitors in DED, as modeled further below.

Lastly, addition of an artificial tears eye drop produced the anticipated immediate increase in tear height and reduction in osmolality, followed by a return to pre-drop steady-state over approximately 2 hours, consistent with clinical observations on the duration of artificial tear efficacy in humans [25]. The multiphasic effect on tear film osmolality is the consequence of relatively rapid transepithelial osmosis compared to re-equilibration of tear fluid ionic contents.

### Dry eye disease simulations

DED is characterized by reduced tear fluid volume and increased osmolality as a consequence of reduced lacrimal gland secretion (aqueous-deficient dry eye) and/or increased evaporation (evaporative dry eye) [2]. However, the distinction between canonical aqueous-deficient and evaporative DED is blurred, with both defects generally occurring together [26]. Based on reported data [27], we chose J_v_ ^lac^ and J_v_^evap^ parameters to model mild/moderative DED to give tear height of 5.4 µm, 27% less than that of 7.4 µm in the healthy eye, and an osmolality of 342 mOsm, 10 mOsm greater than that of 332 mOsm in healthy eyes.

Under dry eye conditions an approximately 2-fold reduction in evaporation fully normalized osmolality to 332 mOsm (Fig. 7A), yet produced little increase in tear height, in agreement with expectations from the simulation done in Fig. 6. In contrast, decreased nasolacrimal drainage or increased lacrimal secretion produced substantial gains in tear fluid height, accompanied by modest reductions in osmolality. Therefore, the model predicts that therapies aimed at reducing evaporation, such as protective goggles or lipid-replenishing eye drops, primarily target hyperosmolarity with limited impact on normalizing tear volume, and that interventions such as punctal plugs to reduce drainage or stimulation of lacrimal secretion primarily normalize tear volume but offer incomplete relief from osmotic stress. Our computations suggest that optimal dry eye treatment requires a combination of evaporation-lowering and volume-enhancing approaches to normalize both tear fluid osmolality and volume. That said, our simulations capture only immediate, short-term shifts; clinically, osmolality and tear thickness are intertwined via inflammation and evaporation dynamics, so that sustained improvement in one typically correlates with at least some recovery in the other over extended periods [2].

**Figure 7.**
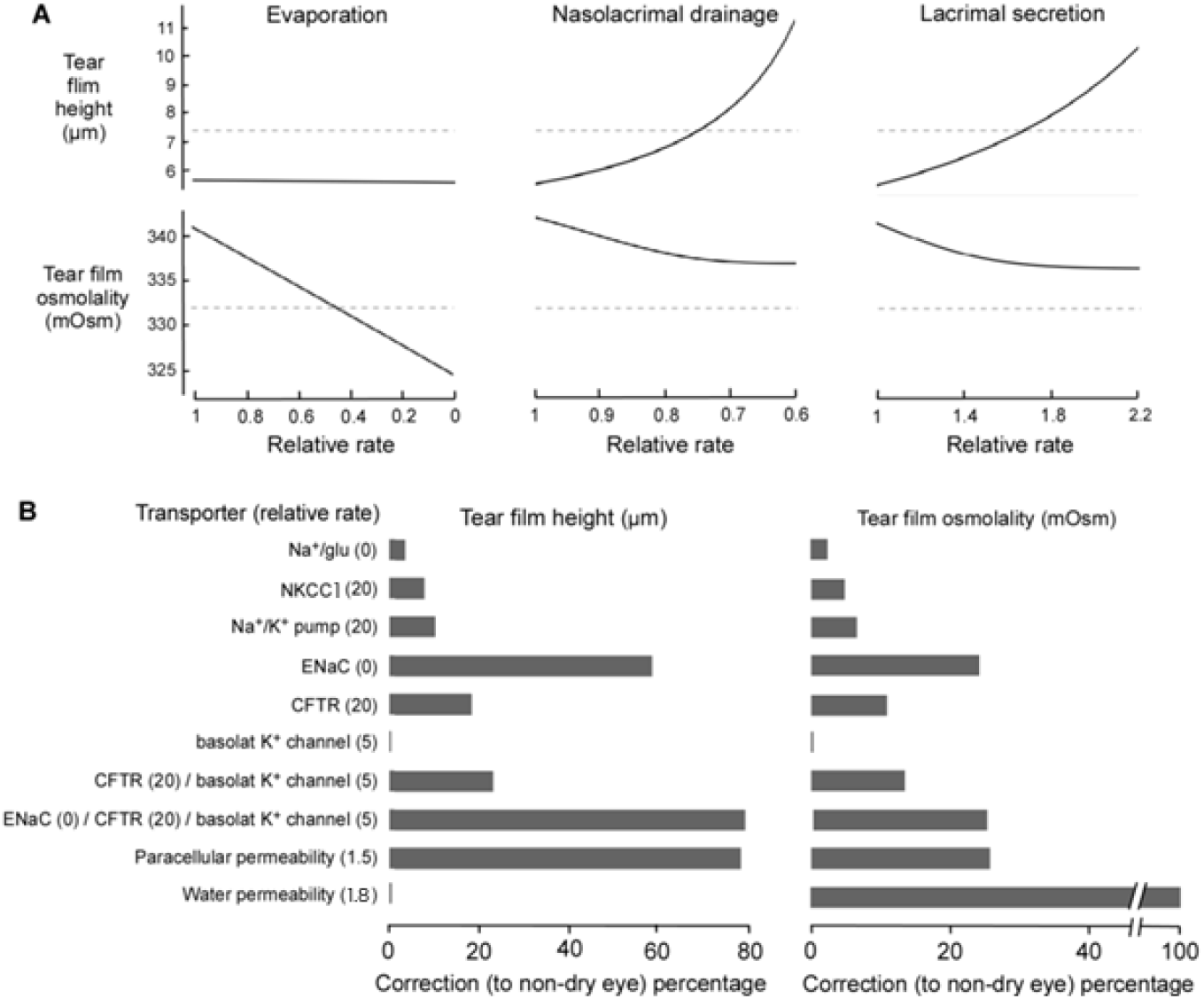
Effects of modifying ocular or epithelial transport parameters on tear fluid height and osmolality in DED. DED defined by a 2.15-fold increase in evaporation and 2.5-fold reduction in lacrimal secretion compared to baseline parameters. A. Effects of reduced evaporation (left), reduced nasolacrimal drainage (middle) and increased lacrimal secretion (right) on tear film height (top) and osmolality (bottom). Dashed horizontal grey lines indicate baseline tear height (7.4 µm) and osmolality (332 mOsm). B. Influence of altered epithelial transport parameters (alone or in indicated combinations) on tear film height (left) and osmolality (right). For each parameter (except for water and paracellular permeabilities), a fold-scaling factor, shown in parentheses, was set to give near-maximal responses.

Lastly, in the context of dry eye we modeled the impacts of modifying epithelial transport parameters, including those targeted by drugs, on tear fluid height and osmolality (Fig. 7B). Corrections are expressed as percentages compared with the healthy, non-dry eye state, with 100% denoting full normalization. ENaC inhibition and increased paracellular permeability were each predicted to increase tear volume while producing only a modest, ∼20%, recovery of tear osmolality. A significant observation was the synergistic action of co-activating CFTR and basolateral K□ channels, which yielded greater improvements than individual activations alone. This synergy arises because CFTR activation alone induces membrane depolarization, reducing the electrochemical driving force for Cl□ efflux; pairing CFTR activation with basolateral K□ channel stimulation maintains hyperpolarization and consequent CFTR Cl□ efflux and fluid secretion. Notably, the triple combination of ENaC inhibition, CFTR activation, and basolateral K^+^-channel stimulation afforded near-maximal normalization of tear height, offering synergistic pro-secretory and anti-absorptive actions. None of the aforementioned modulators predict notable improvements in tear fluid osmolality. Changes to NKCC1, the Na^+^/glucose cotransporter, or the Na□/K□ pump, produced only minor responses.

Effective normalization of tear film height, and to a lesser extent osmolality, was produced by a small, 1.5-fold, increase in paracellular ion permeability. Most remarkably, effective normalization of tear film osmolality was achieved by a 1.8-fold increase in epithelial cell osmotic water permeability, surpassing the corrections achieved by other interventions. Indeed, increasing water permeability by only 1.3-or 1.5-fold produced 50% and 75%, respectively, normalizations of tear fluid osmolality (not shown in figure). These computations predict paracellular ion permeability and cellular water permeability as novel targets for treatment of the abnormal tear film in dry eye (see Discussion).

## Discussion

The model herein was developed to investigate the role of the ocular surface epithelium in regulation of the composition and volume of tear fluid overlying the epithelial layer and to discover novel targets to treat DED. The modeling builds on prior work from our laboratory in the development of the OSPD method to study ion transport at the ocular surface and its applications to the mouse [28], rabbit [29] and human [21] eyes, the development of an electrokinetic model of ocular surface epithelium [17], and the discovery and advancement of dry eye therapeutics targeting ocular surface epithelia [16,30,31]. The modeling accounted for the roles of lacrimal gland secretion, tear fluid evaporation, and nasolacrimal drainage in tear fluid homeostasis, using published evidence on their quantitative contributions and mechanisms. Notably, the modeling also included the principal apical and basolateral membrane transporters that participate in epithelial solute and water transport, accounting for their physiological mechanisms including electrochemical driving forces and transporter saturation, as well as paracellular ion transport. Model parameters were selected systematically, based in part on new OSPD experiments, and the model explicitly required mass balance of each transported component individually in both the tear and cytoplasmic compartments.

The modeling herein builds on prior models of epithelial transport and tear film dynamics, yet differs in several key respects. Prior epithelial transport models have generally used thermodynamic descriptions of coupled ion and water movement across epithelia, representing the epithelium as a single barrier with lumped permeabilities and driving forces [32,33]. Prior tear film models captured the balance of lacrimal secretion, evaporation, drainage, and conjunctival exchange at the level of global fluid and solute mass balance, but without resolving individual epithelial transport mechanisms [34–36]. Here, we combine and extend these approaches using explicit descriptions of each of the major solute and water transport pathways in ocular surface cell membranes and the paracellular pathway, and account explicitly for movements of individual solutes and water in the cytoplasmic and tear fluid compartments, taking into account lacrimal secretion, evaporation, and nasolacrimal drainage. In doing so, we retain the strengths of prior models, including rigorous conservation laws and realistic tear film boundary conditions, while adding mechanistic, transporter-level resolution and enforcing solute-specific mass balance. This integrated formulation made it possible to quantitatively relate changes in individual epithelial transport processes to tear film height and osmolality, permitting direct modeling of therapeutic responses and dry eye phenotypes in a way that was not achievable in earlier models.

Our model provided unique information to guide drug development for treatment of DED. The model supported the development of pro-absorptive and anti-secretory drug candidates targeting the ocular surface epithelium, including ENaC inhibitors [14,37] and CFTR/K□ channel activators [38,39]. However, the predicted efficacy of these drug candidates was modest, particularly for their action on normalization of tear fluid osmolality. Consistent with these predictions, ENaC inhibition with the topical agent P-321 did not meet its primary outcome in a phase 2 clinical trial [4], which might reflect its pharmacodynamics resulting in non-sustained and partial ENaC inhibition. The model predicted more robust effects by the use of pro-absorptive and anti-secretory drugs in combination, supporting the efficacy of a dual anti-absorptive and pro-secretion drug mix. The modeling also predicted the lack of efficacy of targeting the Na□/glucose cotransporter SGLT2, the Na□/K□/2Cl□ cotransporter NKCC1, and the 3Na□/2K□ ATPase, for which approved drug modulators exist (canagliflozin [40], bumetanide [41] and ouabain [42], respectively). A novel and remarkable finding was the predicted high efficacy of increasing water permeability of the ocular surface epithelium, which permits rapid water secretion from the blood to the tear film, thereby normalizing tear fluid osmolality, the principal driver in DED. Pharmacological modulators of the expression of the ocular surface aquaporin AQP5 may thus be therapeutically efficacious, as has been reported for other aquaporins outside of the eye [43], or perhaps aquaporin gene delivery, as has been reported for aquaporins in other epithelia [44,45].

Another interesting and unanticipated result was the effect of modulating paracellular ion transport on tear fluid properties, in which relatively modest, physiologically achievable increases in tight-junctional Na□/K□/Cl□ permeability produced disproportionately large increases in tear fluid volume without producing excessive tear fluid or cytoplasmic hyperosmolality. The corneal surface is lined by a non-keratinized stratified squamous epithelium with 5-7 cell layers, whereas the conjunctival surface is lined by a more heterogeneous, non-keratinized stratified columnar epithelium of 3-5 cell layers with mucus-producing goblet cells, blood vessels, and weaker cell-cell contacts [46]. Tight junction integrity in these epithelia is dependent on claudin and occludin proteins [47], and studies in non-ocular epithelia have identified reversible modulators of these complexes — including calcium chelating agents, medium-chain fatty acids, indomethacin, and inflammatory cytokines — that transiently increase paracellular permeability [48,49]. Targeting the intrinsically leakier conjunctival epithelium with tight junction modulators may therefore normalize the tear fluid with minimal adverse effects on corneal transparency.

The effects of CFTR activation on normalization of tear film properties, with or without basolateral K^+^-channel activation, were relatively modest (Fig. 7B) and are consistent with the limited efficacy seen for CFTR activator therapy in mouse dry eye models. The question arises why CFTR activation in ocular surface epithelia produces only modest changes in tear fluid properties, whereas CFTR activation in the intestine, where transcellular Cl^−^ and fluid secretion are mechanistically similar (apical CFTR with basolateral NKCC1 and K^+^ channels) [50], can produce massive fluid secretion when stimulated by cholera toxin. In ocular surface epithelium CFTR activation (Fig. 6B) was associated with an increase in transepithelial fluid secretion of 2 x 10^-6^ mL cm^-2^ s^-1^, which as predominantly due to increased transcellular secretion. In cholera, fluid secretion can reach 7 x 10^-5^ mL cm^-2^ s^-1^ [51], producing > 1 liter/hour stool water loss [52]. Several differences between intestinal and ocular epithelia likely account for this 35-fold difference in surface area-normalized fluid secretion, including: (i) substantially greater CFTR abundance in intestinal secretory cells than in cornea or conjunctiva [53], (ii) greater secretory driving forces with sustained cyclic AMP activation in cholera [54], and (iii) concurrent inhibition of Na^+^ absorption in cholera [55].

Several limitations of our model are noted, though we do not believe that they alter the principal predictions regarding effects of epithelial transport modulators or tear fluid homeostasis in DED. (i) The ocular surface epithelium was modeled as a homogeneous cell monolayer, whereas the actual ocular surface contains distinct, multi-layered corneal and conjunctival epithelia. Although reported data suggest that these cell types have similar absorptive and secretory transport mechanisms [56], they likely differ to some extent and such differences could impact computed results. Though a straightforward extension of the model here can include both epithelial cell types and their 3-dimensional organization, there is insufficient experimental data to specify the many additional required parameters. (ii) Neither blink-cycle kinematics and surface pressures nor menisci/lid-wiper regions were included in the model [34,36], which would add a significant level of complexity and uncertainty in tear fluid dynamics. (iii) Although the model included the major fluid-transporting components, Na□, K□, Cl□, glucose and water, it did not include bicarbonate, calcium, magnesium, or phosphate, mainly because insufficient information exists about their transporting mechanisms in ocular surface epithelia as well as information about cytoplasmic and tear fluid buffer capacity and carbonic anhydrase kinetics. Inclusion of divalent cations and phosphate is unlikely to alter results because of their low concentrations [57]. (iv) Lastly, our model used a robust set of evidence-based parameters for the mouse eye, with extrapolations from the human eye where appropriate such as in modeling dry eye physiology. Use of all-human parameters, albeit not currently available, may alter the absolute magnitudes of the effects of various modeled maneuvers but is unlikely to alter the principal conclusions.

Further refinements of the tear fluid model are possible building on the modeling framework established herein. As mentioned, extensions of the model could include additional ionic species, distinct corneal and conjunctival cell types, blink-driven fluid pumping dynamics, meniscus effects, and dynamic description of lipid layer breakup. Parameter selection for the human eye may be informative as well using the limited set of OSPD measurements made on human subjects [21]. It may be interesting to incorporate the epithelial model here into spatially resolved tear-film or eyelid-motion models to capture the non-uniform thinning and breakup observed in DED [34,36,58]. Lastly, modeling the actions of inflammatory mediators on epithelial transport and tight-junction integrity would expand the scope of clinically relevant hypotheses and therapeutic strategies that can be investigated [59].

## Experimental methods

### Animals

BALB/c mice (female and male, age 8-16 weeks) were bred at the UCSF Laboratory Animal Resource Center. Protocols were approved by the University of California, San Francisco, Committee on Animal Research and were in compliance with the ARVO Statement for the Use of Animals in Ophthalmic and Vision Research.

### Solutions and reagents

All perfusion solutions were isosmolar to mouse serum (310 mOsm) as measured by freezing point-depression osmometry (Precision Systems, Inc., Natick, MA). Solutions were prepared in one-liter batches with pH titrated to 7.4. All chemicals were purchased from Sigma-Aldrich (St. Louis, MO, USA). Solution compositions are listed in Table 2.

**Table 2.** Composition (in mM) of perfusate solutions

| **Solution** | **Na<sup>+</sup>** | **Cl<sup>-</sup>** | **K<sup>+</sup>** | **sucrose** | **choline<sup>+</sup>** | **gluconate<sup>-</sup>** |
| --- | --- | --- | --- | --- | --- | --- |
| baseline | 137 | 182.4 | 48 | 0 | 0 | 0 |
| 0 Cl<sup>-</sup> | 142 | 0 | 48 | 0 | 0 | 184.9 |
| 0 Na<sup>+</sup> | 0 | 182.4 | 48 | 0 | 137 | 0 |
| 0 K<sup>+</sup> | 142.6 | 182.4 | 0 | 0 | 42.4 | 0 |
| non-ionic | 0 | 0 | 5.6 | 306 | 0 | 0 |

All solutions also contained (in mM): 1 D-glucose, 2.8 phosphate, 1 Ca<sup>2+</sup>, 0.5 Mg<sup>2+</sup>

### OSPD measurements

As described previously [16,17,21,60], mice were anesthetized with isoflurane and kept at 37°C on a heating pad. Open-circuit transepithelial potential differences were recorded during continuous perfusion of the ocular surface with test solutions delivered at 5-10 mL/min. The measuring Ag/AgCl electrode (3 M KCl agar bridge) contacted the perfusion catheter and the reference electrode was grounded using an agar-bridge, 23-gauge butterfly needle inserted subcutaneously in the back. Signals were recorded using high-impedance amplifier/voltmeter interfaced to a computer. Solutions were gravity-fed for 1-3 min until a stable OSPD signal was obtained. Reported OSPD values are the means over the final 10 s of each perfusion. 24 trials were conducted across 9 experimental protocols using specified combinations of perfusate solutions (Table 2) and inhibitors as described in Results.

## Supplementary Information

## Supplementary Methods

### Steady-state solution of tear fluid model

The tear fluid model was independently solved in the steady-state to validate the results of the kinetic computation indicated in Fig. 2 of the main text. The 12 non-linear equations (Eqns. S1-S12) were solved to compute the 12 variables: cytoplasmic solute concentrations [Na^+^]_cell_, [K ^+^]_cell_, [Cl^-^]_cell_, [glu]_cell_; apical/basolateral potentials ψ_ap_, ψ_bl_; layer heights h_cell_, h_tear_; and tear solute concentrations [Na^+^]_tear_, [K^+^]_tear_, [Cl^-^]_tear_, [glu]_tear_. The equations enforce open-circuit ionic current balance (Eqs. S1, S2), and net zero solute and water fluxes into and out of the cytoplasmic (Eqs. S3-S7) and tear fluid (Eqs. S8-S12) compartments.

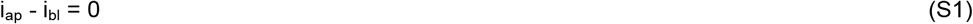

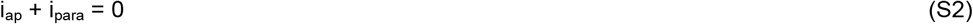

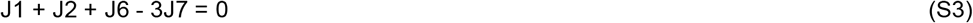

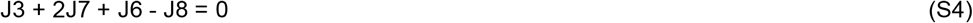

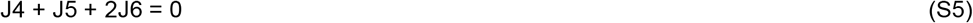

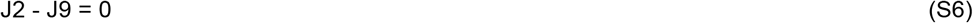

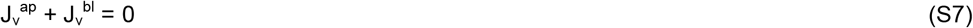

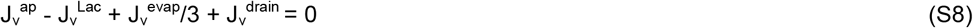

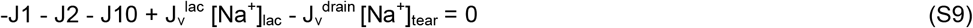

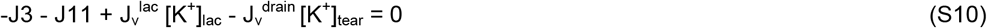

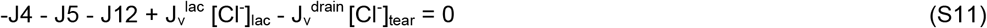

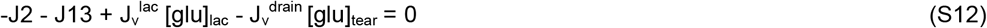

Transporter fluxes, ion currents, and water flow are as defined in Eqns. 1-16 in the main text, and lacrimal secretion, tear film evaporation, and nasolacrimal drainage are as defined in Supplementary Figs. S1-S3. In Eq. S8 J_v_^evap^ is divided by 3 to account for evaporation occurring only over the exposed ocular surface in the mouse eye. This ratio was computed by comparing images of exposed ocular surface area [1] with total tear surface area, A_tear_. A_tear_ was estimated by scaling the human tear film surface area (cornea and conjunctiva) by the human-to-mouse tear-volume ratio [2,3] and correcting for the thinner human tear film (∼ 50% of that in mice) [4].

The non-linear equation system was solved with SciPy’s bounded least_squares [5], enforcing physiological variable bounds and using a high evaluation cap (max_nfev=10^5^) with stringent stopping tolerances (ftol=10^-15^, xtol=10^-15^, gtol=10^-15^) and a minimum acceptable residual of 10^-12^.

### Tear film kinetics following topical application of artificial tears

Addition of a droplet of artificial tears of volume V_drop_ and osmolality Osm_drop_ was modeled. The immediate post-instillation tear osmolality or solute X concentration is, [X](t+Δt) = ([X](t) h_tear_(t) + [X]_drop_ Δh_tear_) ÷ (h_tear_(t) + Δh_tear_), where A_tear_ is as estimated above and Δh_tear_ = V_drop_ / A_tear_. In Fig. 6, the added fluid had a composition similar to mouse blood and within the range of standard artificial tear formulations (osmolality 325 mOsm, Na^+^ 112 mM, K^+^ 3.2 mM, Cl^−^ 84 mM) [6].

### Tear film evaporation, lacrimal gland secretion, and nasolacrimal drainage Tear film evaporation

Net tear film evaporation rate, J_v_^evap^, is given by,

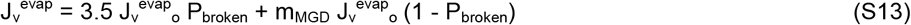

where J_v_^evap^_o_is the baseline evaporation rate for freshly blinked eyes (13.6 g/cm^2^/s [7]), P_broken_is fractional area of tear breakup, m_MGD_is the relative evaporation from intact tear areas representing lipid deficiency (1.0 for healthy eye, 1.6 for dry eye [7]), and the factor 3.5 accounts for relative evaporation from broken vs. intact tear area [8,9].

Equation S13 describes the area-weighted average evaporation rate across the ocular surface, accounting for broken and intact regions of the tear film. Immediately after a blink the tear film is at its thickest and most uniform with P_broken_near zero and evaporation across the intact lipid layer at the baseline rate of J_v_^evap^_o_m_MGD_. During the inter-blink period the film thins unevenly, increasing P_broken_and creating localized breakup areas where the tear film ruptures, exposing the underlying epithelium.

P_broken_is taken as a logistic function of tear film height based on data relating breakup percentage to tear film height [10] (Supplementary Fig. S1, top),

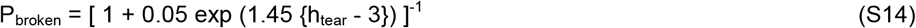

In healthy eyes ∼4% of the tear film breaks up over a typical inter-blink interval, compared to ∼23% in dry eye due to reduced film height and stability [10]. Dry eyes also have higher baseline evaporation in intact areas due to lipid deficiency, as quantified by m_MGD_. Supplementary Fig. S1 (bottom) shows J_v_^evap^ as a function of h_tear_in healthy and dry eye.

### Lacrimal secretion

Lacrimal fluid composition varies with lacrimal gland secretion rate, with higher secretion rates producing fluid relatively isosmolar to plasma, with increasing K^+^ and Cl^−^ and reduced Na^+^ concentrations. For the modeling here, we interpolate osmolality and ion concentrations between basal and stimulated secretion rates using an exponential function,

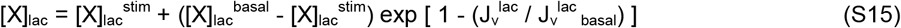

where J_v_^lac^ _basal_is basal secretion rate and X is osmolality, Na^+^, K^+^ or Cl^-^. Supplementary Table S4 lists stimulated vs. basal osmolality and ion concentrations, and Supplementary Fig. S2 shows tear fluid osmolality and ion concentrations as function of lacrimal secretion rate J_v_^lac^._Basal_Glucose concentration is taken to be independent of secretion rate, at 7.0 mM. As needed, a non-ionic, impermeant osmolyte is included to match the target osmolality with that computed from ionic concentrations. The basal lacrimal secretion rate in mice J ^lac^ is 1.23 x 10^-6^ cm/s [11]. Basal lacrimal gland fluid concentrations were taken from Walcott et al. [12] and stimulated concentrations were extrapolated from studies in rabbits [13–15] and humans [16].

### Nasolacrimal drainage

Nasolacrimal drainage removes excess tear fluid from the ocular surface through the nasolacrimal duct. The drainage rate J_v_^drain^ is modeled as a function of tear height (h_tear_),

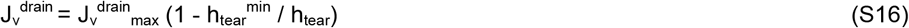

where J_v_^drain^_max_is the maximum possible drainage rate, and h_tear_^min^ is the minimum tear height below which drainage ceases (Supplementary Fig. S3). This function was adapted from Cerretani and Radke [17], which provides a simplified formulation of the more detailed canalicular fluid transport model developed by Zhu and Chauhan [18]. Drainage rate (J_v_^drain^_max_) has not been measured in mice, but was specified here to give a baseline steady-state tear height of 7.4 µm [19], giving J_v_^drain^_max_= 3.2 x 10^-6^ cm/s. h_tear_^min^ for mice was taken as 3.0 µm based on drainage measurements and modeling in human dry eye [18].

## Supplementary Tables

**Supplementary Table S1.**
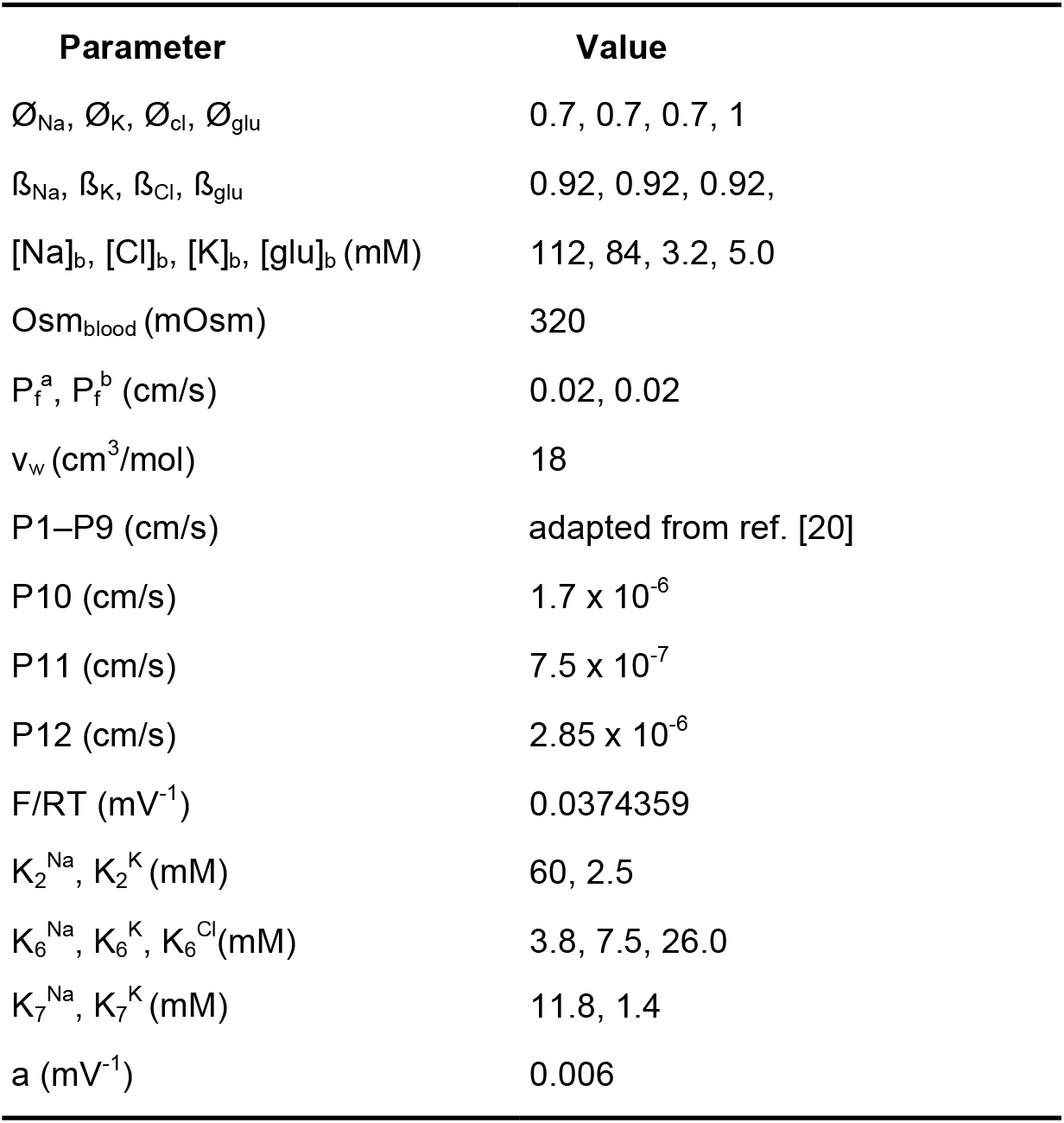
Model parameters (for baseline, non-dry eye)

| Parameter | Value |
| --- | --- |
| $\emptyset_{\text{Na}}, \emptyset_{\text{K}}, \emptyset_{\text{Cl}}, \emptyset_{\text{glu}}$ | 0.7, 0.7, 0.7, 1 |
| $\beta_{\text{Na}}, \beta_{\text{K}}, \beta_{\text{Cl}}, \beta_{\text{glu}}$ | 0.92, 0.92, 0.92, |
| $[\text{Na}]_{\text{b}}, [\text{Cl}]_{\text{b}}, [\text{K}]_{\text{b}}, [\text{glu}]_{\text{b}}$ (mM) | 112, 84, 3.2, 5.0 |
| $\text{Osm}_{\text{blood}}$ (mOsm) | 320 |
| $P_{\text{f}}^{\text{a}}, P_{\text{f}}^{\text{b}}$ (cm/s) | 0.02, 0.02 |
| $v_{\text{w}}$ (cm<sup>3</sup>/mol) | 18 |
| P1–P9 (cm/s) | adapted from ref. [20] |
| P10 (cm/s) | $1.7 \times 10^{-6}$ |
| P11 (cm/s) | $7.5 \times 10^{-7}$ |
| P12 (cm/s) | $2.85 \times 10^{-6}$ |
| $F/RT$ (mV<sup>-1</sup>) | 0.0374359 |
| $K_2^{\text{Na}}, K_2^{\text{K}}$ (mM) | 60, 2.5 |
| $K_6^{\text{Na}}, K_6^{\text{K}}, K_6^{\text{Cl}}$ (mM) | 3.8, 7.5, 26.0 |
| $K_7^{\text{Na}}, K_7^{\text{K}}$ (mM) | 11.8, 1.4 |
| $a$ (mV<sup>-1</sup>) | 0.006 |

**Supplementary Table S2.**
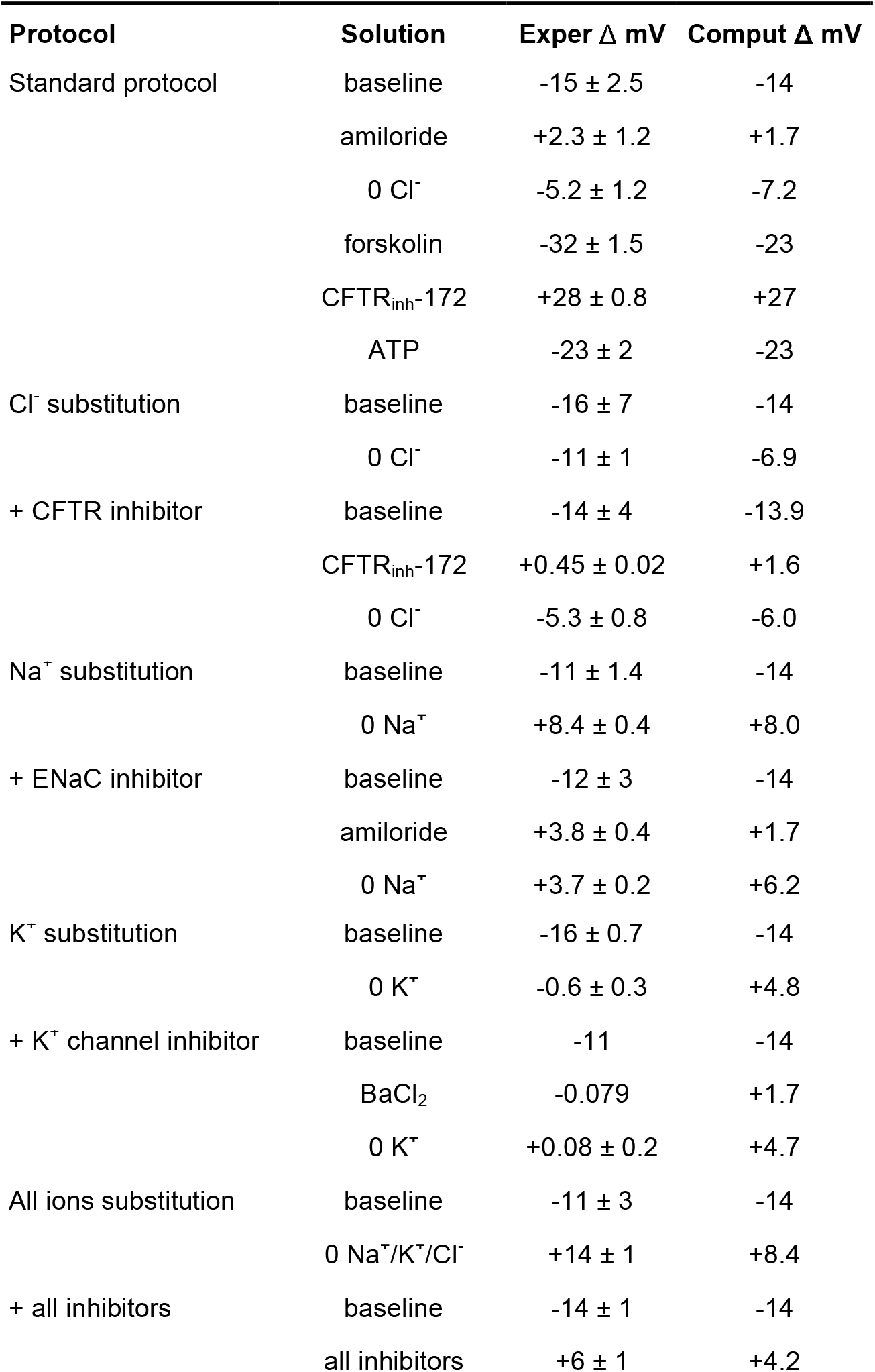
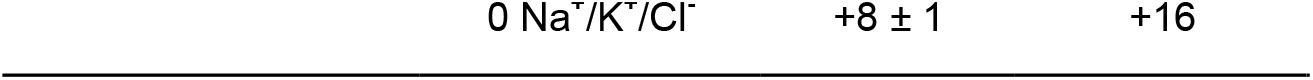
Experimental and computed OSPD results. Changes in OSPD (Δ mV) for indicated experimental maneuvers as in Fig. 3. Δ mV values are mean ± S.E.M. for up to 4 measurements. Computed Δ mV determined using model parameters listed in Table S1.

| Protocol | Solution | Exper $\Delta$ mV | Comput $\Delta$ mV |
| --- | --- | --- | --- |
| Standard protocol | baseline | -15 ± 2.5 | -14 |
| | amiloride | +2.3 ± 1.2 | +1.7 |
| | 0 Cl<sup>-</sup> | -5.2 ± 1.2 | -7.2 |
| | forskolin | -32 ± 1.5 | -23 |
| | CFTR<sub>inh</sub>-172 | +28 ± 0.8 | +27 |
| | ATP | -23 ± 2 | -23 |
| Cl<sup>-</sup> substitution | baseline | -16 ± 7 | -14 |
| | 0 Cl<sup>-</sup> | -11 ± 1 | -6.9 |
| + CFTR inhibitor | baseline | -14 ± 4 | -13.9 |
| | CFTR<sub>inh</sub>-172 | +0.45 ± 0.02 | +1.6 |
| | 0 Cl<sup>-</sup> | -5.3 ± 0.8 | -6.0 |
| Na<sup>+</sup> substitution | baseline | -11 ± 1.4 | -14 |
| | 0 Na<sup>+</sup> | +8.4 ± 0.4 | +8.0 |
| + ENaC inhibitor | baseline | -12 ± 3 | -14 |
| | amiloride | +3.8 ± 0.4 | +1.7 |
| | 0 Na<sup>+</sup> | +3.7 ± 0.2 | +6.2 |
| K<sup>+</sup> substitution | baseline | -16 ± 0.7 | -14 |
| | 0 K<sup>+</sup> | -0.6 ± 0.3 | +4.8 |
| + K<sup>+</sup> channel inhibitor | baseline | -11 | -14 |
|  | BaCl<sub>2</sub> | -0.079 | +1.7 |
| | 0 K<sup>+</sup> | +0.08 ± 0.2 | +4.7 |
| All ions substitution | baseline | -11 ± 3 | -14 |
| | 0 Na<sup>+</sup>/K<sup>+</sup>/Cl<sup>-</sup> | +14 ± 1 | +8.4 |
| + all inhibitors | baseline | -14 ± 1 | -14 |
| | all inhibitors | +6 ± 1 | +4.2 |

**Supplementary Table S3.** Baseline computed and reported tear film parameters for healthy (non-dry eye) mice. Note the difference in computed vs. literature [Cl^-^]_tear_reflects the fact that the model does not contain bicarbonate, with the elevated [Cl^-^]_tear_required for electroneutrality. Literature values taken from refs [19–24].

| Parameter | Computed | Literature |
| --- | --- | --- |
| OSPD (mV) | -12.1 | -13.9 |
| [Na<sup>+</sup>]<sub>cell</sub> (mM) | 19 | 20 |
| [K<sup>+</sup>]<sub>cell</sub> (mM) | 80 | 75 |
| [Cl<sup>-</sup>]<sub>cell</sub> (mM) | 31 | 33 |
| [glu]<sub>cell</sub> (mM) | 7.6 | 20 |
| [Na<sup>+</sup>]<sub>tear</sub> (mM) | 137 | 139 ± 8 |
| [K<sup>+</sup>]<sub>tear</sub> (mM) | 45 | 48 ± 1 |
| [Cl<sup>-</sup>]<sub>tear</sub> (mM) | 161 | 127 ± 4 |
| [glu]<sub>tear</sub> (mM) | 0.734 | 1.05 ± 0.36 |
| Osm<sub>tear</sub> (mOsm) | 332 | 323-338 |
| h<sub>tear</sub> (μm) | 7.4 | 7.4 |

**Supplementary Table S4.** Tear fluid osmolality and composition benchmarks of basal and stimulated lacrimal gland fluid.

| Parameter | Basal | Stimulated |
| --- | --- | --- |
| Osm<sub>lac</sub> (mOsm) | 345 | 330 |
| [Na<sup>+</sup>]<sub>lac</sub> (mM) | 150 | 144 |
| [K<sup>+</sup>]<sub>lac</sub> (mM) | 38 | 46 |
| [Cl<sup>-</sup>]<sub>lac</sub> (mM) | 155 | 159 |

**Supplementary Table S5:** Steady-state parameters at relative lacrimal secretion rates (J_v_^lac^) of 0.1, 1, and 2-fold baseline. All other parameters held constant.

| Parameter | Relative secretion rate |  |  |
| --- | --- | --- | --- |
|  | 0.1x | 1x | 2x |
| OSPD (mV) | -12 | -12 | -12 |
| $\psi_{\text{ap}}$ (mV) | -43 | -63 | -63 |
| $\psi_{\text{bl}}$ (mV) | -54 | -75 | -75 |
| $[\text{Na}^+]_{\text{cell}}$ (mM) | 16 | 19 | 20 |
| $[\text{K}^+]_{\text{cell}}$ (mM) | 38 | 80 | 82 |
| $[\text{Cl}^-]_{\text{cell}}$ (mM) | 53 | 31 | 30 |
| $[\text{glu}]_{\text{cell}}$ (mM) | 5.5 | 7.6 | 9.5 |
| $[\text{Na}^+]_{\text{tear}}$ (mM) | 165 | 137 | 140 |
| $[\text{K}^+]_{\text{tear}}$ (mM) | 22 | 45 | 48 |
| $[\text{Cl}^-]_{\text{tear}}$ (mM) | 182 | 161 | 163 |
| $[\text{glu}]_{\text{tear}}$ (mM) | 0.24 | 0.73 | 1.40 |
| $\text{Osm}_{\text{tear}}$ (mOsm) | 347 | 332 | 331 |
| $h_{\text{tear}}$ ($\mu\text{m}$) | 4.3 | 7.4 | 113 |
| $J_v^{\text{ap}}$ ($\text{cm/s} \times 10^{-6}$) | -4.8 | -2.2 | -2.0 |
| Transcellular flux ($\text{mOsm/cm/s} \times 10^{-5}$) | -1.7 | -2.5 | -3.9 |
| Paracellular flux ($\text{mOsm/cm/s} \times 10^{-4}$) | 3.1 | 2.3 | 2.4 |
| $J_v^{\text{drain}}$ ($\text{cm/s} \times 10^{-6}$) | 1.0 | 1.9 | 3.1 |

**Supplementary Table S6.**
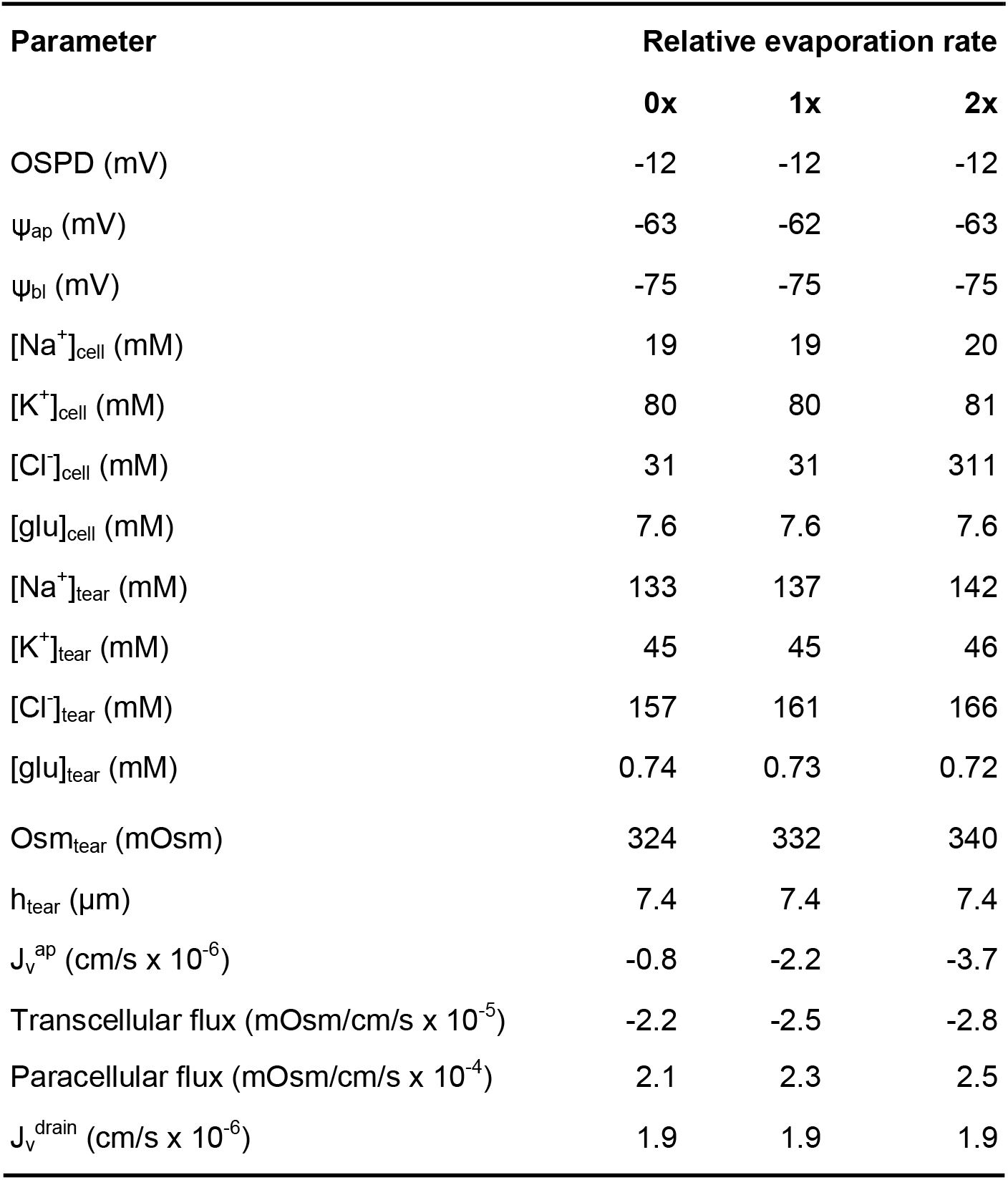
Steady-state parameters at relative evaporation rates of 0, 1, and 2-fold baseline. All other parameters held constant.

| Parameter | Relative evaporation rate |  |  |
| --- | --- | --- | --- |
|  | 0x | 1x | 2x |
| OSPD (mV) | -12 | -12 | -12 |
| $\psi_{ap}$ (mV) | -63 | -62 | -63 |
| $\psi_{bl}$ (mV) | -75 | -75 | -75 |
| $[Na^+]_{cell}$ (mM) | 19 | 19 | 20 |
| $[K^+]_{cell}$ (mM) | 80 | 80 | 81 |
| $[Cl^-]_{cell}$ (mM) | 31 | 31 | 311 |
| $[glu]_{cell}$ (mM) | 7.6 | 7.6 | 7.6 |
| $[Na^+]_{tear}$ (mM) | 133 | 137 | 142 |
| $[K^+]_{tear}$ (mM) | 45 | 45 | 46 |
| $[Cl^-]_{tear}$ (mM) | 157 | 161 | 166 |
| $[glu]_{tear}$ (mM) | 0.74 | 0.73 | 0.72 |
| $Osm_{tear}$ (mOsm) | 324 | 332 | 340 |
| $h_{tear}$ ($\mu m$) | 7.4 | 7.4 | 7.4 |
| $J_v^{ap}$ (cm/s $\times 10^{-6}$) | -0.8 | -2.2 | -3.7 |
| Transcellular flux (mOsm/cm/s $\times 10^{-5}$) | -2.2 | -2.5 | -2.8 |
| Paracellular flux (mOsm/cm/s $\times 10^{-4}$) | 2.1 | 2.3 | 2.5 |
| $J_v^{drain}$ (cm/s $\times 10^{-6}$) | 1.9 | 1.9 | 1.9 |

**Supplementary Table S7.**
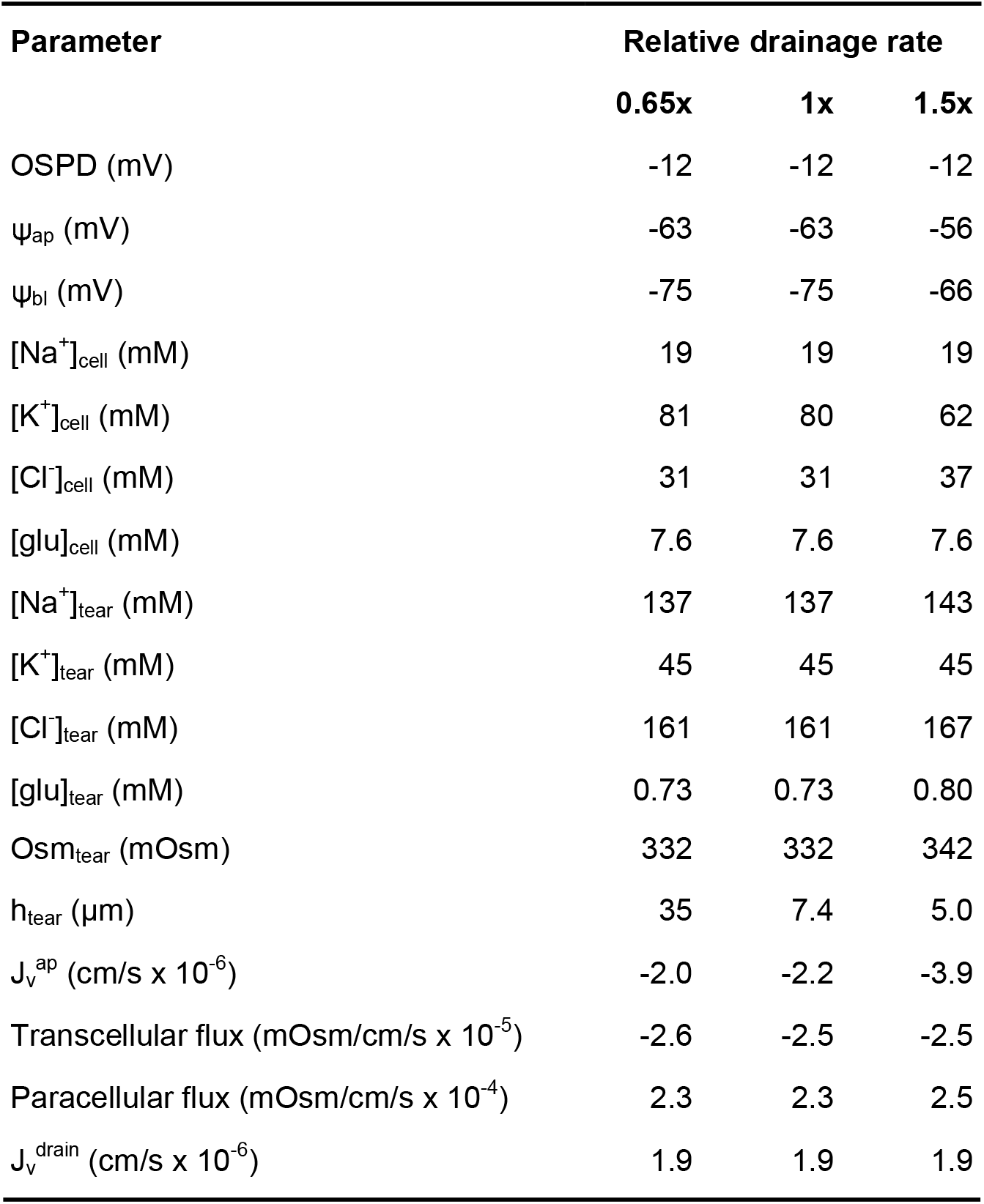
Steady-state parameters at relative nasolacrimal drainage rates of 0.65, 1, and 1.5-fold baseline. All other parameters held constant.

| Parameter | Relative drainage rate |  |  |
| --- | --- | --- | --- |
|  | 0.65x | 1x | 1.5x |
| OSPD (mV) | -12 | -12 | -12 |
| $\psi_{ap}$ (mV) | -63 | -63 | -56 |
| $\psi_{bl}$ (mV) | -75 | -75 | -66 |
| $[Na^+]_{cell}$ (mM) | 19 | 19 | 19 |
| $[K^+]_{cell}$ (mM) | 81 | 80 | 62 |
| $[Cl^-]_{cell}$ (mM) | 31 | 31 | 37 |
| $[glu]_{cell}$ (mM) | 7.6 | 7.6 | 7.6 |
| $[Na^+]_{tear}$ (mM) | 137 | 137 | 143 |
| $[K^+]_{tear}$ (mM) | 45 | 45 | 45 |
| $[Cl^-]_{tear}$ (mM) | 161 | 161 | 167 |
| $[glu]_{tear}$ (mM) | 0.73 | 0.73 | 0.80 |
| Osm<sub>tear</sub> (mOsm) | 332 | 332 | 342 |
| $h_{tear}$ ($\mu m$) | 35 | 7.4 | 5.0 |
| $J_v^{ap}$ (cm/s x $10^{-6}$) | -2.0 | -2.2 | -3.9 |
| Transcellular flux (mOsm/cm/s x $10^{-5}$) | -2.6 | -2.5 | -2.5 |
| Paracellular flux (mOsm/cm/s x $10^{-4}$) | 2.3 | 2.3 | 2.5 |
| $J_v^{drain}$ (cm/s x $10^{-6}$) | 1.9 | 1.9 | 1.9 |

## Supplementary Figures

**Supplementary Figure S1.**
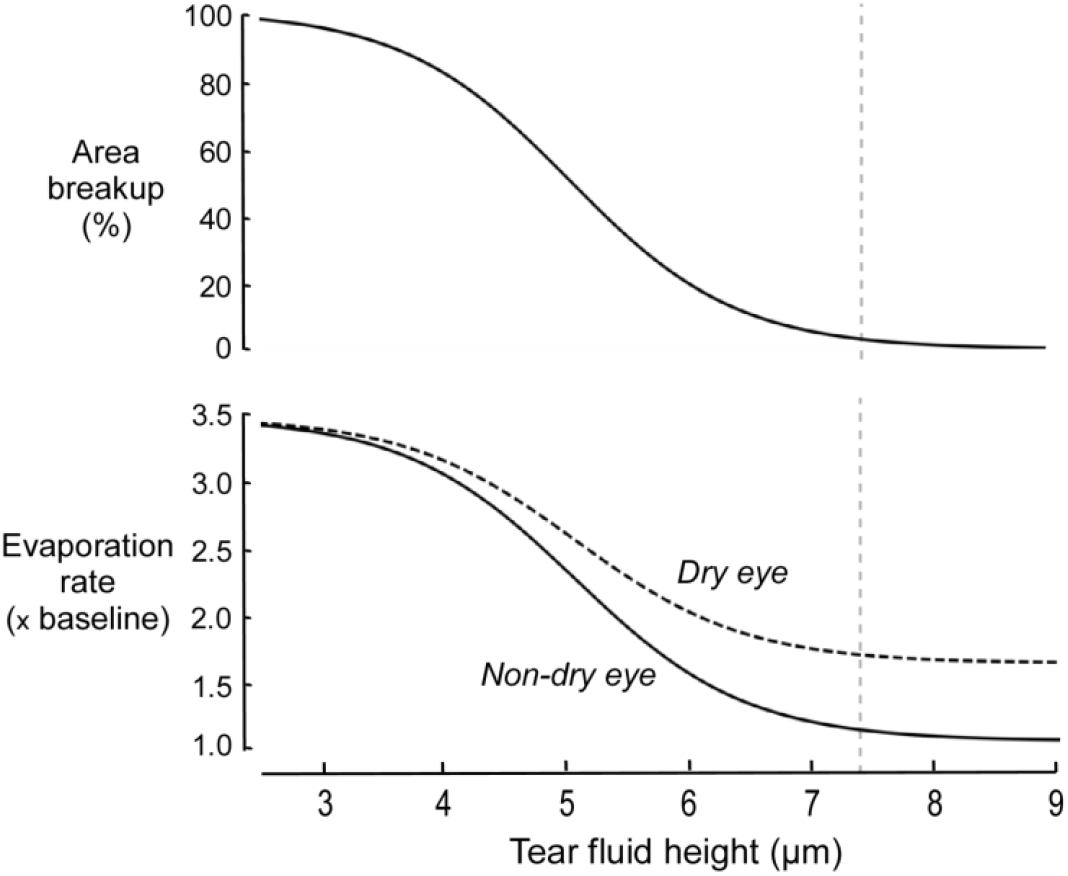
Dependence of percentage tear breakup area (P_broken_) (top) and average evaporation rate (bottom) on tear fluid height. Evaporation rate shown in dry eye and healthy non-dry eye. Vertical line drawn at basal h_tear_(7.4 µm).

**Supplementary Figure S2.**
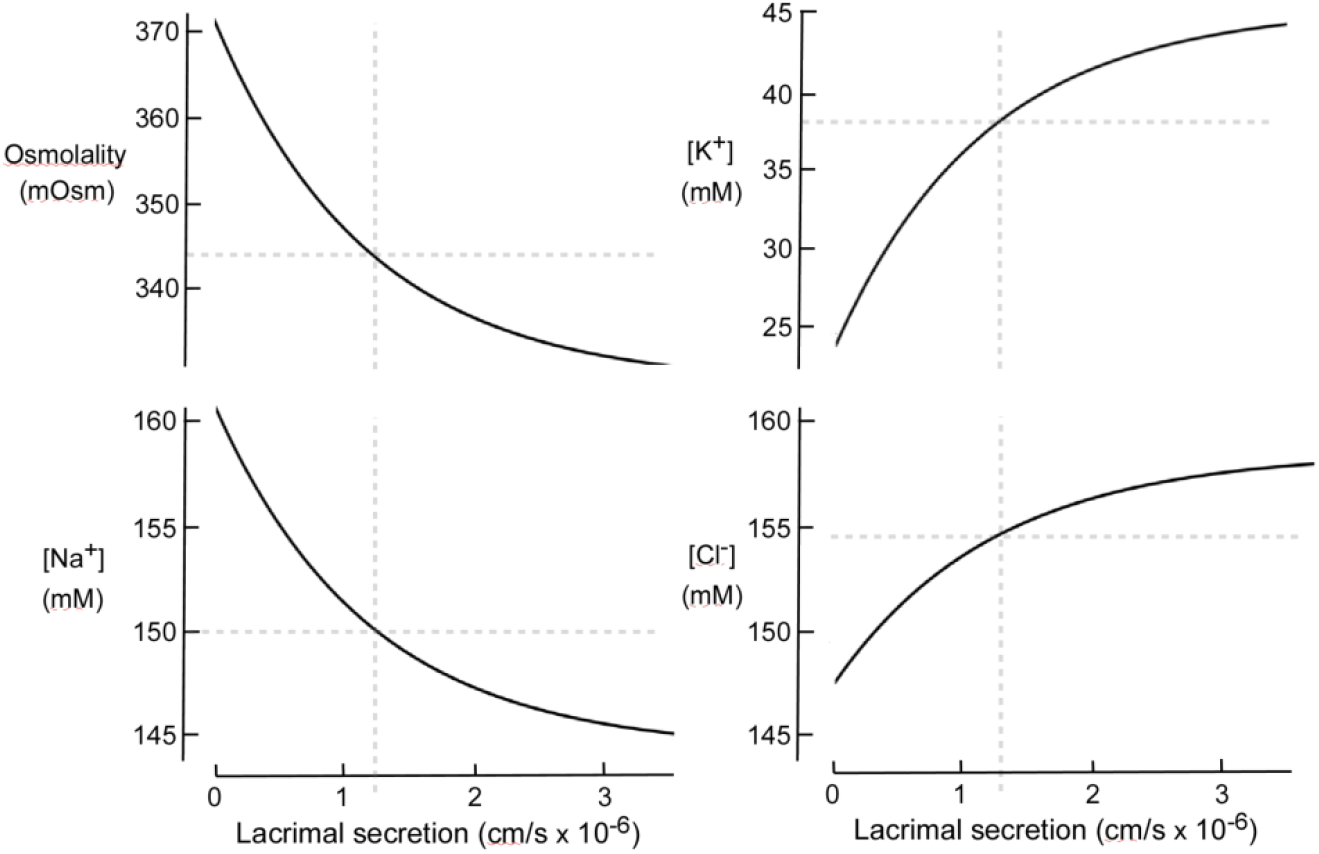
Tear fluid osmolality and ion concentrations as a function of lacrimal secretion rate J_v_^lac^. Dashed horizontal grey lines indicate basal osmolality or concentration (Osm_lac_, [X]_lac_); dashed vertical grey lines at basal J_v_^lac^ (1.23 x 10^-6^ cm/s).

**Supplementary Figure S3.**
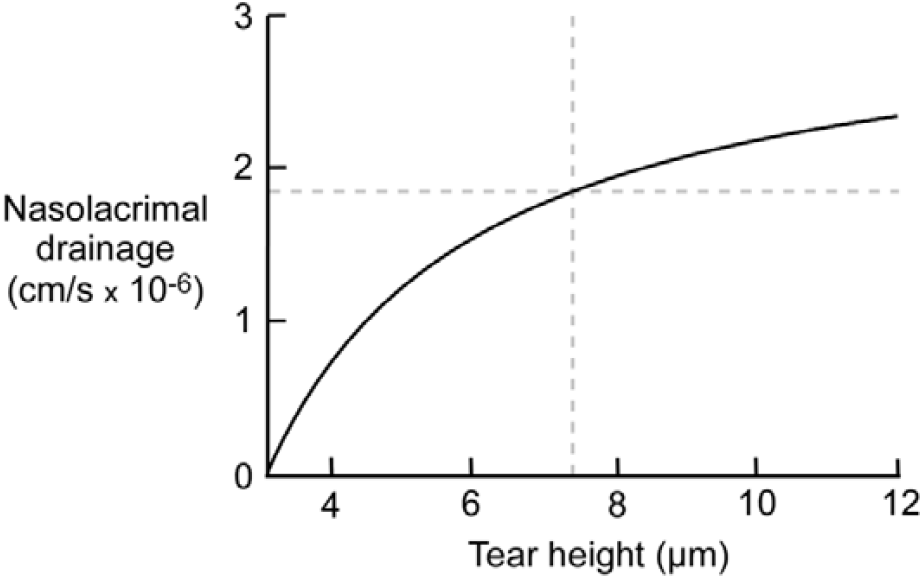
Nasolacrimal drainage rate as a function of tear film height. Vertical line drawn at baseline h_tear_(7.4 µm).

